# The Degradation-promoting Roles of Deubiquitinases Ubp6 and Ubp3 in Cytosolic and ER Protein Quality Control

**DOI:** 10.1101/2020.01.22.915793

**Authors:** Hongyi Wu, Davis T.W. Ng, Ian Cheong, Paul Matsudaira

**Affiliations:** Temasek Life Sciences Laboratory, 1 Research Link, National University of Singapore, Singapore 117604; Department of Biological Sciences, National University of Singapore, Block S3 #05-01, 16 Science Drive 4, Singapore 117558; Mechanobiology Institute, National University of Singapore, T-Lab #10-01, 5A Engineering Drive 1, Singapore 117411

## Abstract

The quality control of intracellular proteins is achieved by degrading misfolded proteins which cannot be refolded by molecular chaperones. In eukaryotes, such degradation is handled primarily by the ubiquitin-proteasome system. However, it remains unclear whether and how protein quality control deploys various deubiquitinases. To address this question, we screened deletions or mutation of the 20 deubiquitinase genes in *Saccharomyces cerevisiae* and discovered that almost half of the mutations slowed the removal of misfolded proteins whereas none of the remaining mutations accelerated this process significantly. Further characterization revealed that Ubp6 maintains the level of free ubiquitin to promote the elimination of misfolded cytosolic proteins, while Ubp3 supports the degradation of misfolded cytosolic and ER luminal proteins by different mechanisms.

## Introduction

Protein quality control (QC) pathways operate in all compartments of eukaryotic cells to eliminate misfolded proteins, the accumulation of which correlates with various age-onset diseases (Choe et al., 2016; Kaushik and Cuervo, 2015; Klabonski et al., 2016). In cytosolic QC (CytoQC), chaperones bind misfolded proteins to inhibit aggregation and assist with refolding (Park et al., 2007). Substrates which fail to refold, such as Ste6*c and ΔssPrA, are degraded by the ubiquitin-proteasome system (UPS) (Figure S1 and Heck et al., 2010). Since many chaperones shuttle between the cytosol and the nucleus, misfolded cytosolic proteins can thus be ferried into the nucleus to be degraded by the nuclear UPS (Park et al., 2013; Prasad et al., 2018). Aggregates of cytosolic proteins can be re-solubilized by chaperones and degraded via the UPS or directly cleared by macro-autophagy (Yu et al., 2018). Similarly, in the endoplasmic reticulum (ER), proteins which misfold in their luminal, transmembrane, or cytosolic domains are engaged by respective ERQC-L, -M and -C machineries (Vashist and Ng, 2004), and are retro-translocated into the cytosol for degradation by the UPS (Hampton and Sommer, 2012). The model substrates of ERQC include CPY*, Sec61-2 and Ste6* (Figure S1).

The UPS, which is responsible for degrading the majority of misfolded proteins, consists of the proteasomes and enzymes which catalyze protein ubiquitination, namely the ubiquitin-activating enzyme (E1), -conjugating enzyme (E2) and -ligating enzyme (E3) (Finley et al., 2012). Additionally, deubiquitinases (DUbs) such as Ubp6 and Doa4 in *Saccharomyces cerevisiae* (budding yeast) recycle ubiquitin from ubiquitinated proteins and ubiquitin-fusion proteins (Figure S2 and Amerik et al., 1997; Chernova et al., 2003; Hanna et al., 2003; Ozkaynak et al., 1987; Swaminathan et al., 1999; Tobias and Varshavsky, 1991). Deubiquitination by various DUbs also regulates different processes such as transcription, translation, signal transduction and vesicle transport (Reyes-Turcu et al., 2009). For instance, Ubp3 in yeast deubiquitinates Sec23 to facilitate protein transport between the ER and Golgi by COPII vesicles (Cohen et al., 2003a; Cohen et al., 2003b).

Although DUbs function in a variety of cellular activities, little is known about the spectrum of DUbs involved in QC or the exact roles of a few DUbs implicated in QC pathways, such as Ubp3 and Ubp6. Ubp3 supports CytoQC under heat stress by suppressing the conjugation of lysine 63 (K63)-linked ubiquitin chains on misfolded proteins and facilitating K48-linkage (Fang et al., 2014; Fang et al., 2016; Silva et al., 2015) but its function under the physiological temperature or in other QC pathways is unknown (Oling et al., 2014). Ubp6 was proposed to delay QC because deleting *UBP6* reduced the steady-state abundance of some proteins (Boselli et al., 2017; Nielsen et al., 2014; Torres et al., 2010). This hypothesis, however, lacks support from direct assays of degradation kinetics (Dephoure et al., 2014).

To resolve the roles of DUbs in QC, we screened deletions or mutation of all DUb genes in *S. cerevisiae* and quantified their effects on CytoQC and ERQC. We found that half of the deletions decelerate QC whereas the other half have no significant effect. Interestingly, *Δubp6*, which was previously suggested to accelerate QC, delays CytoQC by reducing the level of free ubiquitin, but leaves ERQC unaffected. In contrast, *Δubp3* delays ERQC by compromising the transport between ER and Golgi, and also slows the degradation of a subset of CytoQC substrates by a yet uncharacterized mechanism. These findings demonstrate that the Dubs Ubp6 and Ubp3 support different QC pathways by distinct ways.

## Results

### A reverse genetic screen identified DUbs that support QC degradation

We screened all 20 DUbs in *S. cerevisiae* (Figure S2) by measuring the ability of gene deletion or hypomorphic mutation strains to degrade the CytoQC substrate Ste6*c and ERQC substrate CPY* (Guerriero et al., 2013; Prasad et al., 2012; Tran, 2019; Vashist and Ng, 2004). In wild-type (WT), Ste6*c was rapidly degraded with only 30% of the substrate remaining at 12 min post-labeling (Figure 1A). By contrast, the elimination of Ste6*c by CytoQC was significantly slower in *rpn11*^*S119F*^, *Δubp6, Δubp3, Δubp8, Δubp10* and *Δdoa4* (with over 47% of Ste6*c remaining) and moderately slower in *Δubp2, Δubp14, Δotu2* and *Δubp1* (with over 41% remaining) (Figure 1A and S3A). Degradation was slightly faster in *Δubp13* and *Δubp11* (with 20% and 23% remained) but no further acceleration was observed in the *Δubp11Δubp13* double deletion strain (Figure 1A and S4). The remaining 9 single mutants degraded Ste6*c at WT kinetics (Figure 1A, S3A and S4). As for ERQC, *rpn11*^*S119F*^ and *Δubp3* delayed the degradation of CPY* (with over 76% of CPY* remaining compared to 44% in WT) whereas the remaining mutants, including several which delayed CytoQC (*e.g. Δubp6*), eliminated CPY* at WT kinetics (Figure 1B and S3B). Thus, *rpn11*^*S119F*^, *Δubp6* and *Δubp3* impair CytoQC most severely while *rpn11*^*S119F*^ and *Δubp3* also compromise ERQC. The functions of Ubp6 and Ubp3 in CytoQC and ERQC were further explored.

**Figure 1.**
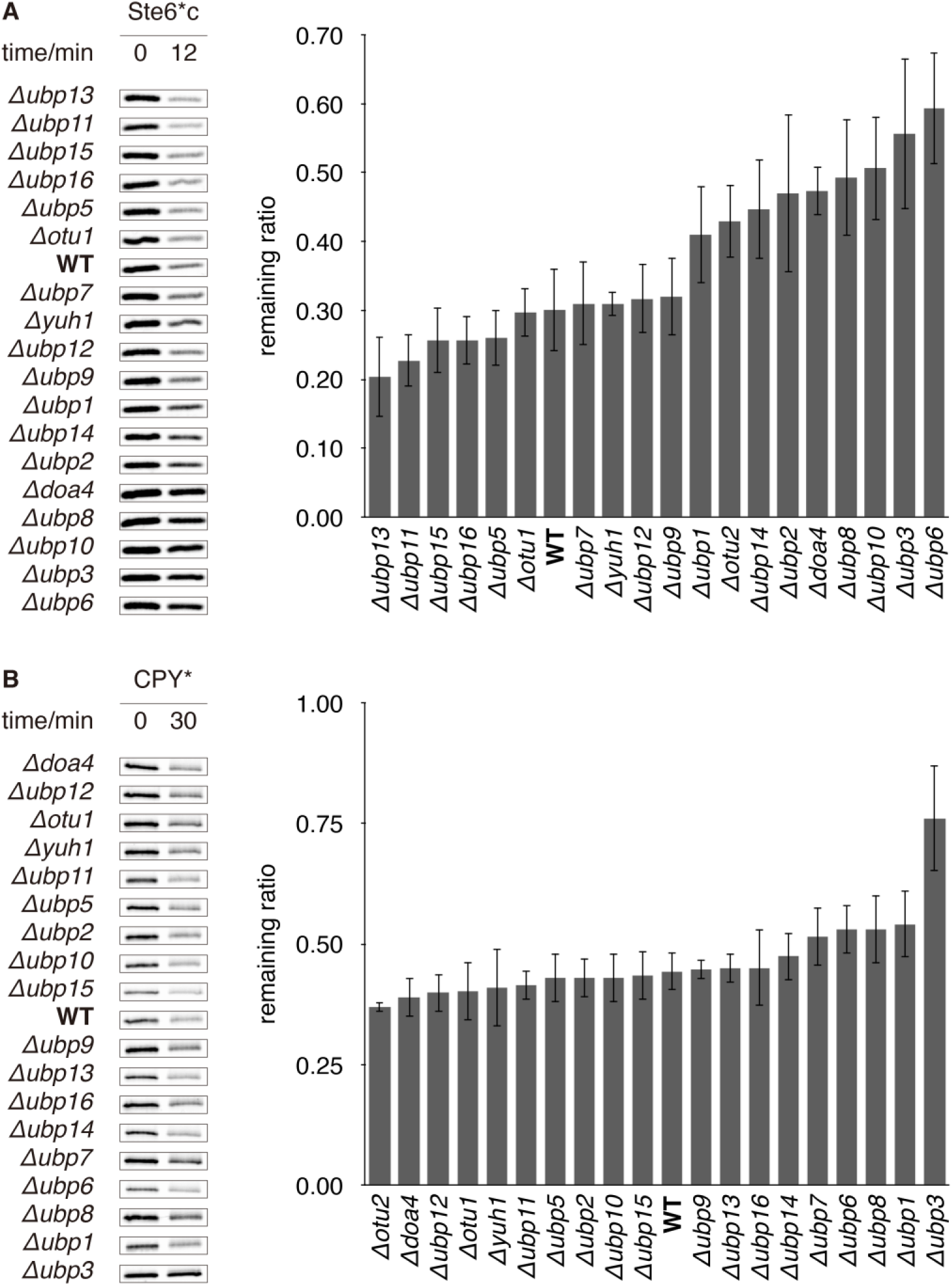
QC is delayed in various DUb deletion strains. (**A**) Degradation of Ste6*c, a misfolded cytosolic protein, and (**B**) degradation of CPY*, a misfolded ER protein in WT versus DUb deletion strains. Cells that express the substrates Ste6*c or CPY* were pulse-labelled with radioisotopic amino acids and sampled at the indicated time-points. The substrates were then immunoprecipitated, resolved by SDS-PAGE, and exposed to storage phosphor screens. Experiments in this study were performed thrice at 30°C unless otherwise stated. Left: representative gel images. Right: quantification of replicate experiments. The remaining ratio of Ste6*c and CPY* was calculated for each strain as the ratio between the remaining abundance at 12 min and 30 min respectively to the initial abundance (at 0 min). Error bars show standard deviations (s.d.). t-tests were performed between mutants and WT. If p-value < 0.05, an asterisk (*) is deposited to the top.

### Ubp6 promotes CytoQC

Ubp6 is a peripheral subunit of the proteasome which recycles ubiquitin from substrates prior to proteolysis (Aufderheide et al., 2015; Bashore et al., 2015; Hanna et al., 2006; Hanna et al., 2003). Although *UBP6* deletion had been suggested to enhance QC (Boselli et al., 2017; Nielsen et al., 2014; Torres et al., 2010), it in fact compromised the degradation of CytoQC substrate Ste6*c and ΔssPrA (Figure 1A, 2A and 2B) and the defect can be complemented by re-expression of Ubp6 from centromeric plasmid (data not shown). Meanwhile, it did not delay or accelerate the clearance of any ERQC substrate, CPY*, Ste6* or Sec61-2 (Figure 1B and 2C - E). In no instance was a QC pathway accelerated.

Because other DUbs that promote QC such as Rpn11, Doa4 and Ubp14 (Figure 1A) are also required for degrading folded proteins by non-QC pathways (Figure S3D and Amerik et al., 1997; Swaminathan et al., 1999), we examined the role of Ubp6 in the degradation of two folded proteins, Stp1 and Deg1-Ura3. Stp1 is a transcription factor, whose uncleaved cytosolic (immature, i) and cleaved nuclear (mature, m) forms (Figure 2F) are degraded rapidly by the UPS (Tumusiime et al., 2011). Deg1-Ura3 is the fusion of Ura3 to the degradation signal (Deg1) of MATα2 and is localized in the cytosol (Zattas et al., 2016). While the elimination of misfolded proteins requires chaperones to maintain solubility or recruit E3, Stp1 and Deg1-Ura3 are degraded in a chaperone-independent manner, which justifies them as folded substrates (Gowda et al., 2013; Hickey, 2016; Park et al., 2007). In *rpn11*^*S119F*^, which served as a control, the degradation of Deg1-Ura3 was significantly decelerated (Figure S3D) whereas in *Δubp6* both Deg1-Ura3 and Stp1 were degraded at WT kinetics (Figure 2F and G). These results suggest that Ubp6 acts specifically in clearing misfolded proteins by CytoQC.

**Figure 2.**
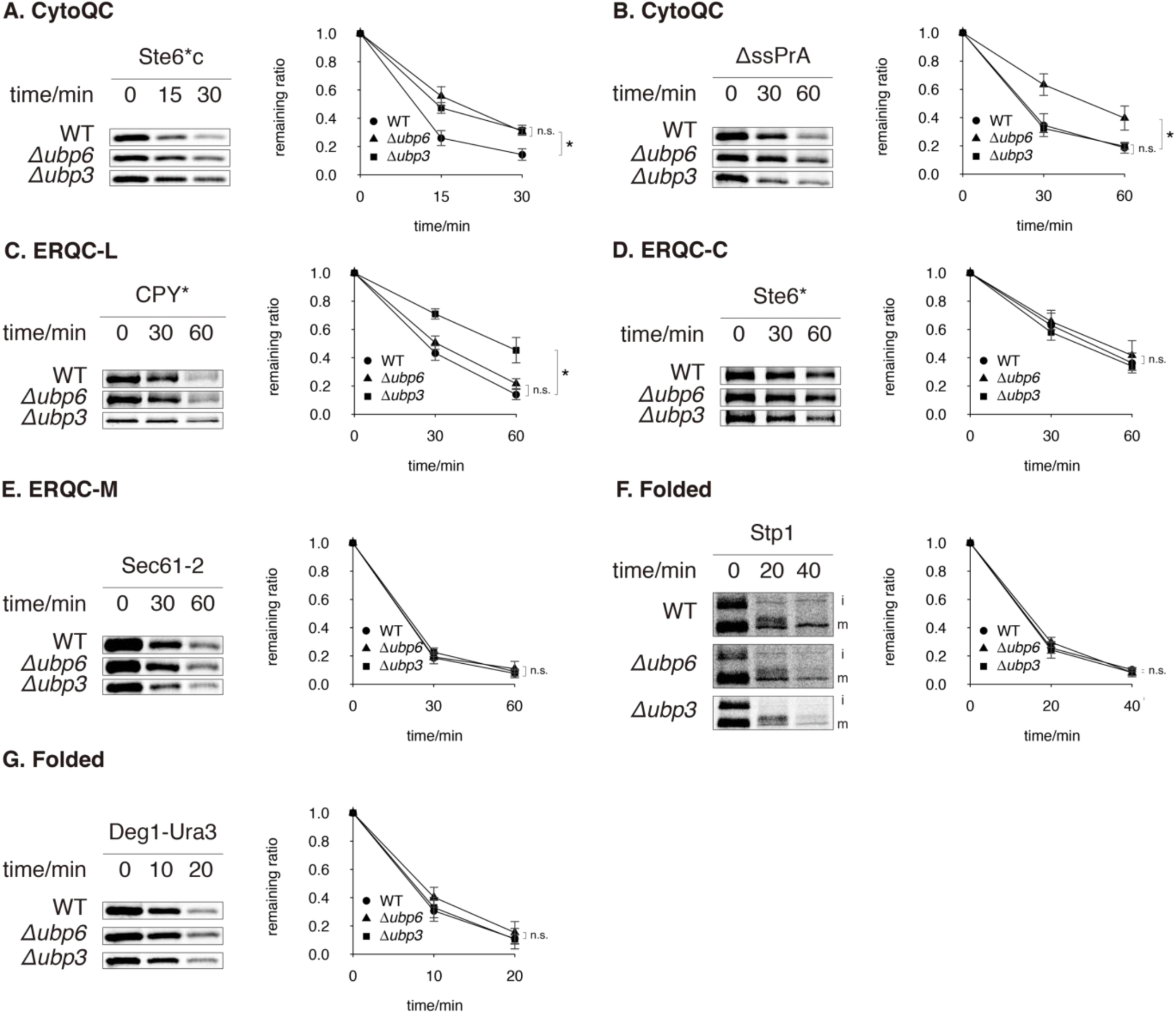
*Δubp6* and *Δubp3* compromise different QC pathways. (**A and B**) Degradation of cytosolic misfolded proteins. (**C - E**) Degradation of proteins that misfold in the lumen, on the cytosolic surface and in the membrane of ER, respectively. (**F and G**) Degradation of folded proteins. Uncleaved (immature) and cleaved (mature) forms of Stp1 are indicated as “i” and “m”, respectively. All substrates were pulsed-labeled and sampled at the indicated time-points. Their remaining ratios were plotted against time. t-tests were performed for each time-point between different curves. If p-value < 0.05 in at least one t-test, an asterisk (*) is indicated, otherwise “n.s.” (non-significant) is shown.

Ubp6 was originally proposed to delay degradation in aneuploid strains (Boselli et al., 2017; Nielsen et al., 2014; Torres et al., 2010) although our results in haploids have suggested otherwise. To test the possibility that Ubp6 functions differently in aneuploid strains where genes exist in aberrant copy numbers or are expressed differentially (Dephoure et al., 2014; Pavelka et al., 2010), we assayed QC in an aneuploid strain with duplicated *chromosome XIII* (*dis XIII*), where Ubp6 had been suggested to suppress QC (Boselli et al., 2017; Nielsen et al., 2014; Torres et al., 2010). In *dis XIII*, as in haploid, *UBP6* deletion significantly compromised CytoQC (Figure 3A and B), moderately compromised ERQC-L (Figure 3C), and had no effect on ERQC-C or ERQC-M (Figure 3D and E). At 25°C, which is more commonly used to maintain aneuploid strains (Torres et al., 2010), the same pattern of phenotypes was observed (data not shown). Furthermore, in an unrelated aneuploid strain possessing duplicated *chromosomes I* and *VIII* (*dis I, VIII*), deleting *UBP6* imposed similar delays in CytoQC but less in ERQC (data not shown). The above evidence proves that Ubp6 is required for efficient CytoQC in both haploid and aneuploid yeast.

**Figure 3.**
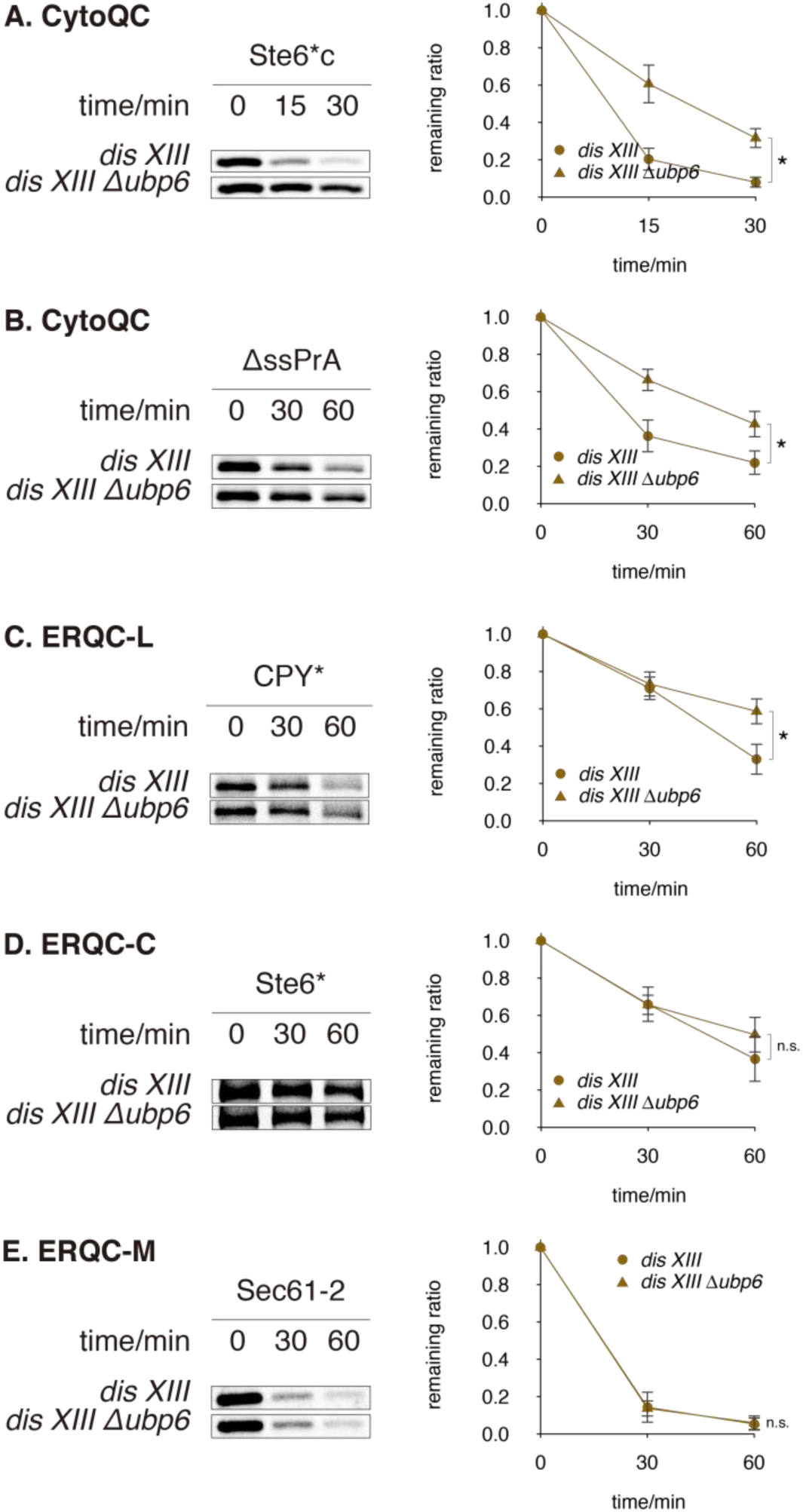
*Δubp6* delays QC in aneuploid strains at 30°C. (**A and B**) Degradation of cytosolic misfolded proteins in *dis XIII* and *dis XIII Δubp6*. (**C - E**) Degradation of ER proteins that misfold in the lumen, on the cytosolic surface and in the membrane, respectively. Substrates were pulsed-chased as in Figure 2.

### Restoring free ubiquitin abundance in *Δubp6* rescues CytoQC

Because Ubp6 is a deubiquitinase, we next examined the levels of ubiquitinated CytoQC substrates in WT and *Δubp6*. In WT, the most abundant species of ubiquitinated Ste6*c or ΔssPrA were tagged with di-ubiquitin chains (Figure 4A). As the chain length increased, ubiquitinated substrates decreased in abundance (Figure 4A and data not shown). In contrast, the abundance of ubiquitinated Ste6*c or ΔssPrA decreased by 50 - 70% in *Δubp6* and the chain lengths were also shorter (Figure 4A and data not shown).

**Figure 4.**
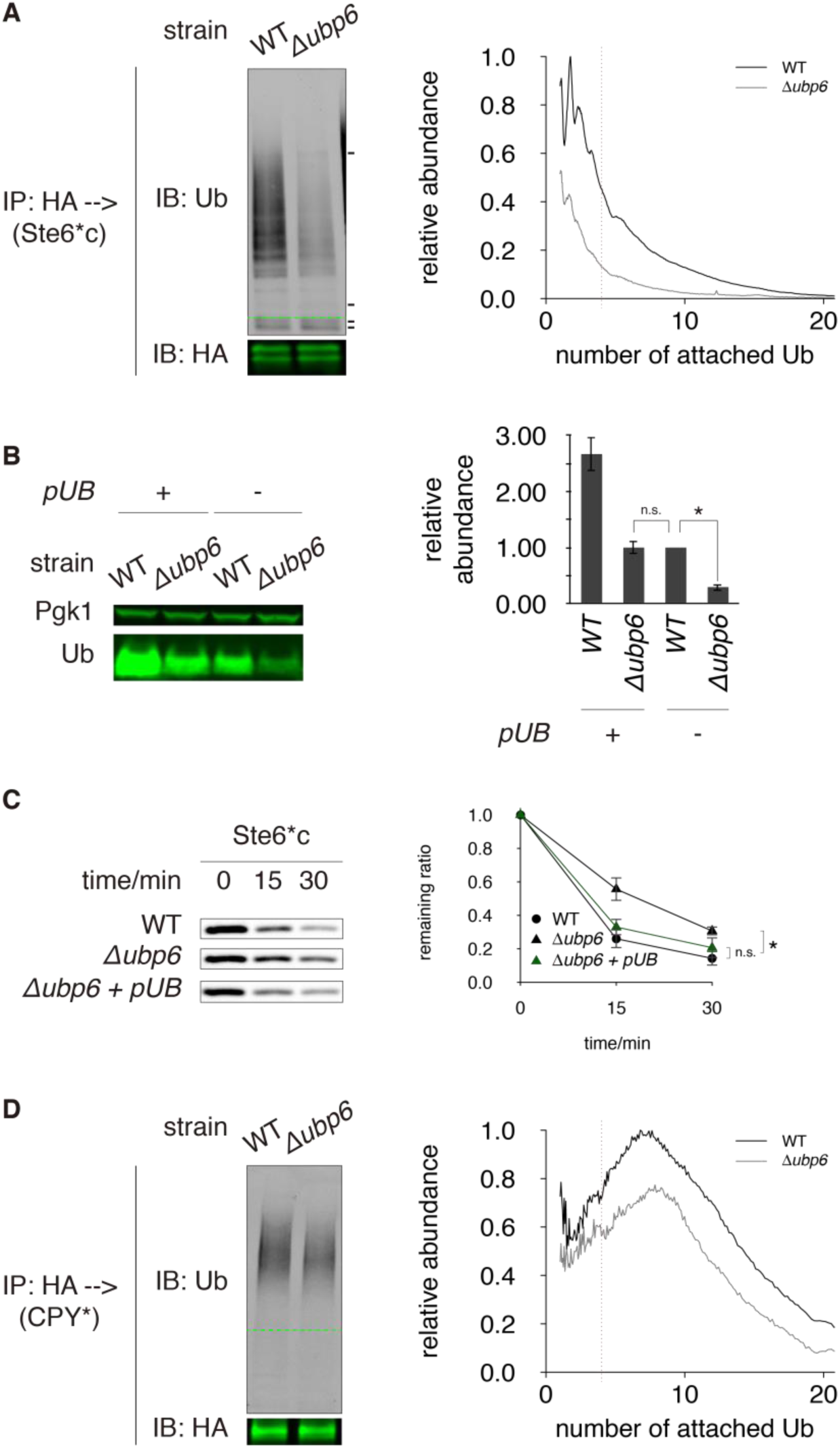
CytoQC is rescued by restoring free ubiquitin abundance in *Δubp6*. (**A**) Ubiquitination of Ste6*c, a cytosolic misfolded protein. Proteins were extracted under non-reducing condition to preserve unconventional ubiquitination on cysteine residues. Ste6*c was immunoprecipitated (IP) using anti-HA affinity matrix, fractionated by SDS-PAGE (non-reducing) and visualized by immunoblotting against ubiquitin (greyscale) and HA (green). Amount of proteins loaded for IP was normalized based on the amounts of non-ubiquitinated proteins. Ubiquitinated proteins are observable as smear and ladder in anti-ubiquitin blots. The positions of non-ubiquitinated substrates are indicated by green dashed lines (---). Non-specific bands, which originate from HA affinity matrix, are indicated by black dashes (**-**). Left: representative blots. Right: profiles of ubiquitination. The number of ubiquitin molecules attached to a ubiquitinated species was calculated from the latter’s molecular weight and plotted along the horizontal axis. The abundance of each species, plotted along the vertical axis, was calculated by normalizing the strength of fluorescent signal to the number of ubiquitin moieties and then to the abundance of non-ubiquitinated substrate. The red dashed vertical line indicates where the ubiquitin chain length is 4, the minimum threshold for high-affinity interaction with proteasome. (**B**) Abundance of free (mono-)ubiquitin in WT and *Δubp6* expressing Ste6*c in the presence and absence of ubiquitin overexpression (*pUB*). Experiments were performed under non-reducing condition as in (A). Pgk1 was probed as a loading control. (**C**) Degradation of Ste6*c in *Δubp6* + *pUB*, shown along with degradation in WT and *Δubp6* (without *pUB*). Ste6*c was pulsed-chased as in Figure 2. (**D**) Ubiquitination of CPY*, a misfolded ER luminal protein. Similar to (A).

Since *Δubp6* exhibited lower ubiquitination levels (Figure 4A), and was known to contain ∼ 60% less free ubiquitin (Hanna et al., 2003 and Figure 4B), we tested whether the free ubiquitin pool limits degradation by CytoQC. When the abundance of free ubiquitin in *Δubp6* was restored to WT level or greater (Figure 4B), the degradation of Ste6*c and ΔssPrA recovered to WT kinetics (Figure 4C and data not shown). In addition, overexpressing ubiquitin in WT almost tripled the abundance of free ubiquitin (Figure 4B) but the kinetics of CytoQC remained the same (Figure S5A). These results indicate that when the free ubiquitin pool decreased in *Δubp6* below WT levels, degradation by CytoQC slowed.

If *Δubp6* decelerates CytoQC by reducing the abundance of free ubiquitin, then why is the degradation kinetics of ERQC not affected by this ubiquitin depletion (Figure 2E - F)? To address this question, we profiled the ubiquitination status of misfolded ER proteins. The ubiquitin chain lengths of substrates in WT peaked at eight molecules for CPY* and at 3 (Figure 4D) and 9 ubiquitin molecules for Sec61-2 (Figure S5B). However, like CytoQC, the abundance of ubiquitinated CPY* and Sec61-2 in *Δubp6* decreased by 50-70% for species tagged with more than 4 ubiquitin molecules and less so for species tagged with 1-3 ubiquitin molecules (Figure 4B and S5B). These profiles showed that although ERQC substrates were degraded at kinetics comparable to WT, they were ubiquitinated to a lesser extent in *Δubp6*, as observed for CytoQC substrates.

### Ubp3 supports CytoQC and ERQC-L

While Ubp3 had been known to support CytoQC under heat stress (Fang et al., 2016), our genetic screen further revealed that it supports both CytoQC and ERQC at the physiological temperature (30°C) (Figure 1A and B). We proceeded to investigate Ubp3’s functions in CytoQC, ERQC and the turnover of folded proteins. Similar to *Δubp6, Δubp*3 delayed the clearance of CytoQC substrate Ste6*c (Figure 1A and 2A), but *Δubp3* did not influence the clearance of ΔssPrA (Figure 2B). ΔssPrA is distinct from Ste6*c in that it localizes to the nucleus and is ubiquitinated by the E3 San1 (Prasad et al., 2010; Prasad et al., 2012; Prasad et al., 2018). However, Δ2GFP, another San1-dependent and nuclear-localized CytoQC substrate (Prasad et al., 2010; Prasad et al., 2018), also depended on Ubp3 for clearance (Figure S6A). These results show that Ubp3 is required for degrading a subset of cytosolic misfolded substrates but the requirement is not determined by substrate localization or E3 preference.

Deleting *UBP3* also decelerated the clearance of misfolded ER luminal protein CPY* (Figure 1B and 2C) but not the integral ER membrane proteins Ste6* or Sec61-2, which contain a mutation in the cytoplasmic or transmembrane portion respectively (Figure 2D and E). To distinguish whether Ubp3 is specifically required by misfolded luminal proteins or any protein with a lesion in the lumen (*i.e.* ERQC-L substrate), we pulse-chased KWW, a membrane protein with a misfolded luminal domain. The degradation of KWW was delayed in *Δubp3* (Figure S6B), which demonstrates that the entire branch of ERQC-L is compromised in *Δubp3*.

Similarly, to determine whether *Δubp3* affects the degradation of folded UPS substrates, we pulse-chased Stp1 and Deg1-Ura3. *Δubp3* did not alter the degradation of these two folded substrates (Figure 2G and H). Therefore, Ubp3 is specifically required by QC pathways, making it similar to Ubp6 and distinct from Rpn11, Doa4 and Ubp14.

### Ubp3 uses distinct mechanisms to support CytoQC and ERQC-L

Under heat stress, Ubp3 promotes CytoQC by exchanging K63-for K48-linkage in ubiquitination so the defect caused by *UBP3* deletion can be surpassed by overexpressing a mutant ubiquitin whose K63 is replaced with arginine (Ub^K63R^) (Fang et al., 2014; Fang et al., 2016; Silva et al., 2015). However, at 30°C when Ub^K63R^ was overexpressed in *Δubp3*, CytoQC and ERQC-L remained slow (Figure 5A and B). This proves that in the absence of heat stress degradation by QC does not depend on K63-ubiquitin linkage removal by Ubp3. Furthermore, the ubiquitination level of CytoQC substrates is the same in *Δubp3* and WT (data not shown) and overexpressing wild-type ubiquitin in *Δubp3* did not rescue CytoQC as in *Δubp6* (Figure 5A), so ubiquitin depletion is not a defect in *Δubp3*.

**Figure 5.**
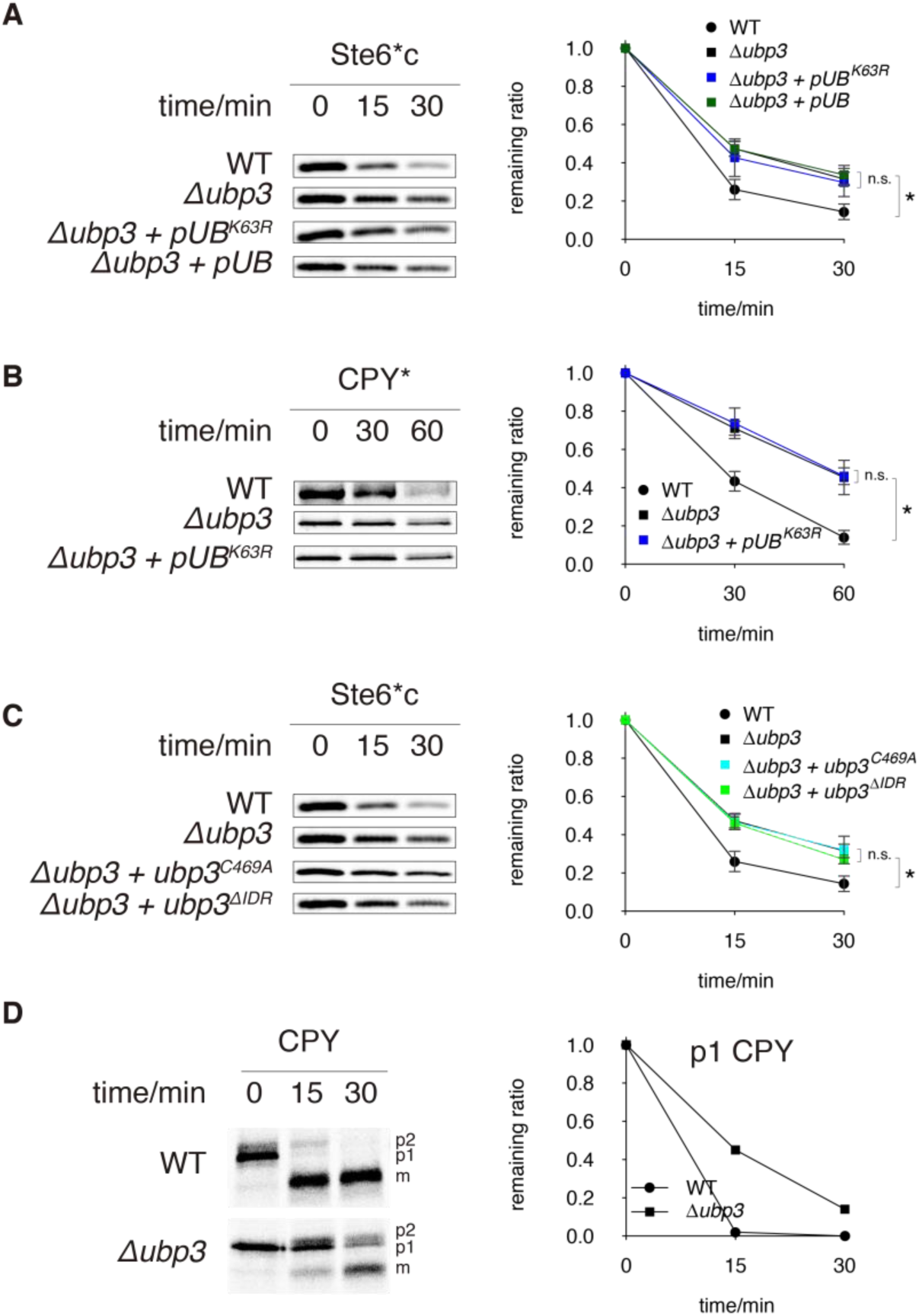
Ubp3 supports QC by distinct mechanisms. (**A**) Degradation of Ste6*c in *Δubp3* + *pUB*^*K63R*^ and *Δubp3* + *pUB*, assayed by pulse-chase as in Figure 2. The degradation in WT and *Δubp3* are shown as controls. (**B**) Degradation of CPY* in *Δubp3* + *pUB*^*K63R*^ assayed by pulse-chase, shown along with degradation in WT and *Δubp3*. (**C**) Degradation of Ste6*c in *Δubp3* + *ubp3*^*C469A*^ assayed by pulse-chase, shown along with degradation in WT and *Δubp3*. (**D**) Maturation of newly synthesized (WT) CPY, assayed by pulse-chase. p1: ER precursor; p2: precursor that has been transported to and modified by the Golgi; m: mature CPY in vacuole. Graph on the right shows the quantification of p1 CPY at different time-points.

*Δubp3* also impairs vesicle transport from the ER to Golgi (Cohen et al., 2003a and Figure 5D). Coincidentally, a delay in ER-to-Golgi transport decelerates ERQC-L but does not affect ERQC-C or -M (Kawaguchi et al., 2010; Taxis et al., 2002; Vashist et al., 2001), which is identical to the phenotype of *Δubp3* (Figure 2C-E). Thus, we reasoned that Ubp3 promotes ERQC-L by facilitating with ER-to-Golgi transport. We further investigated if Ubp3 also facilitates CytoQC by interfering in ER-to-Golgi transport. However, in *sec12-4*, where ER-to-Golgi transport is impaired (Figure S6C), Ste6*c was degraded at WT kinetics (Figure S6D). Therefore, ER-to-Golgi transport is not required by CytoQC. Together, our data demonstrates that *Δubp3* delays CytoQC by a novel mechanism.

To investigate this novel mechanism used by Ubp3, we assayed the roles of its C-terminal DUb domain and a largely disordered region (IDR) at the N-terminus. We generated a catalytically-inactive Ubp3 mutant (Ubp3^C469A^) and a mutant without the IDR (Ubp3^ΔIDR^). These mutants were as stable as WT Ubp3 but were unable to rescue the CytoQC defect in *Δubp3*, demonstrating that the DUb activity and IDR domain are both required for Ubp3 function (Figure 5D).

## Discussion

Protein degradation by QC is delayed when DUbs are overexpressed in cells or included in cell-free reconstitution (Ast et al., 2014; Blount et al., 2012; Liu et al., 2014; Oling et al., 2014; Rumpf and Jentsch, 2006; Zhang et al., 2013). However, in our screen of single deletions or mutation of the 20 DUb genes in *S. cerevisiae*, neither CytoQC nor ERQC was accelerated, and the main phenotype observed was delayed or unaffected degradation of misfolded proteins (Figure 1 and S3A - C). Among DUbs which are required for CytoQC or ERQC, we further characterized Ubp6 and Ubp3 in these pathways.

### Ubp6 promotes CytoQC in many potential ways

In the absence of Ubp6, the level of free ubiquitin is dramatically lower than in WT (Hanna et al., 2003 and Figure 4C), and misfolded cytosolic proteins are less ubiquitinated (Figure 4A) and degraded slower (Figure 1A, 2A and B). As the delay in CytoQC was fully rescued by restoring free ubiquitin levels (Figure 4B and C), the most direct interpretation is that CytoQC is compromised by ubiquitin deficiency. However, other explanations exist as well.

One alternative explanation lies in the proposed competition between Ubp6 and Rpn11, another DUb in the proteasome which removes ubiquitin chains *en bloc* (Guterman and Glickman, 2004; Verma et al., 2002; Yao and Cohen, 2002). Deleting *UBP6* may result in the pre-mature deubiquitination of CytoQC substrates by Rpn11 and dissociation from the proteasomes, giving rise to our observed phenotypes.

Another possibility is that proteasomes become less active in *Δubp6*. According to structural biology analyses, deubiquitination by Ubp6 “lubricates” the translocation of substrates into the proteasome chamber, where proteolysis occurs (Guterman and Glickman, 2004; Verma et al., 2002; Yao and Cohen, 2002). In addition, as a peripheral subunit of the proteasome, Ubp6 can induce conformational change in the proteasome to favor the degradation of certain substrates (Aufderheide et al., 2015; Bashore et al., 2015). If in *Δubp6* the translocation of CytoQC substrates or change in proteasome conformation is hindered, then degradation is slowed and ubiquitinated substrates would accumulate. Nevertheless, if deleting *UBP6* simultaneously delays ubiquitination of CytoQC substrates, the abundance of ubiquitinated substrates could still decrease in *Δubp6*, consistent with what we observed. Similarly, it is possible that certain steps in the pathway is accelerated in *Δubp6* despite an overall delay in CytoQC and still not an acceleration when ubiquitin level is restored (Figure 4C). To reveal if any step in CytoQC is enhanced by *UBP6* deletion will require other techniques such as single-molecule tracking to follow the fates of different populations of misfolded proteins.

### Ubp6 is not required by ERQC or degradation of folded proteins

In contrast to CytoQC, ERQC was not delayed in *Δubp6* even though the substrates were less ubiquitinated (Figure 2C - E, 4D and S5B). This observation supports the notion that the rate-limiting step in ERQC is not ubiquitination but retro-translocation or extraction of proteins from the ER membrane (Guerriero et al., 2016; Nakatsukasa and Kamura, 2016; Wang et al., 2013).

Similarly, the degradation of folded proteins does not require Ubp6. In *Δubp6*, the short-lived folded proteins Stp1-HA and Deg1-Ura3 were degraded at WT rates (Figure 2G and H). Whether the ubiquitination of these folded proteins is affected by *UBP6* deletion remains unclear as we were unable to quantify their ubiquitinated species due to low abundance. Our result is consistent with a previous report that various long-lived proteins in *Δubp6* displayed WT degradation rates (Dephoure et al., 2014). Furthermore, it was reported that short-lived folded proteins Gcn4, Ub-e^K^-K-Trp1 and Ub-e^K^-K-Ura3 had faster turnovers in *Δubp6*, although their accelerated turnover appears like a result of attenuated substrate expression (Hanna et al., 2006). Interestingly, two foreign proteins, Ub-e^K^-K-β-gal and Ub-e^K^-P-β-gal from *Escherichia coli*, rely on Ubp6 for efficient removal (Leggett et al., 2002). This is likely because β-gal in yeast is recognized by chaperones and degraded via CytoQC, even though it can “fold” to be functional (Lee et al., 1996). Together, we can conclude that Ubp6 is not required for the degradation of ERQC or folded substrates but is specific for misfolded cytosolic proteins.

### Ubp3 supports QC at normal temperature

Proteins under heat-stress are decorated with ubiquitin chains of higher content of K63-linkage, as Rsp5, which normally assembles K63-linked ubiquitin chains, becomes a major E3 that catalyzes ubiquitination (Fang et al., 2014; Fang et al., 2016; Silva et al., 2015). Ubp3 was found to associate with Rsp5 under higher temperature to exchange K63-for K48-linkage. Deletion of *UBP3* under the same condition results in slower degradation and accumulation of protein aggregates (Fang et al., 2016; Oling et al., 2014). Our study further revealed that at the physiological temperature, Ubp3 still promotes CytoQC (Figure 1A, 2A and S6A) even though Rsp5 is no longer involved (Fang et al., 2014; Fang et al., 2016; Silva et al., 2015). Interestingly, Ubp3 is required by only a subset of CytoQC substrates, indicating the existence of two CytoQC branches of distinct Ubp3-reliance. Moreover, the promotion by Ubp3 is unrelated to the remodeling of K63 linkage into K48 (Figure 5A) or the level of substrate ubiquitination (data not shown). Currently, we hypothesize that Ubp3 trims ubiquitin chains of certain forked topology, which have been reported to inhibit degradation (Kim et al., 2007). To test this hypothesis, mass-spectrometry must be deployed to determine and compare the topology of ubiquitin chains installed on CytoQC substrates in *Δubp3* and WT.

At 30°C, we also discovered that Ubp3 supports ERQC-L (Figure 2C). Ubp3 utilizes Bre5 as a cofactor to recognize and deubiquitinate Sec23 (Cohen et al., 2003a; Cohen et al., 2003b). This activity is required for ER-to-Golgi transport (Figure 5C), which in turn is implicated in ERQC-L (Taxis et al., 2002; Vashist et al., 2001). However, it is confirmed that ERQC-L substrates *per se* need not undergo ER-to-Golgi transport before degradation (Kawaguchi et al., 2010; Taxis et al., 2002), so the exact mechanism of how ER-to-Golgi transport maintains ERQC-L remains to be unveiled (Taxis et al., 2002).

### The localization and domain organization of DUbs determine their functions in QC

As presented in this article, Ubp6 and Ubp3 use different mechanisms to promote degradation of distinct sets of misfolded proteins (Figure 2). In addition, three other hits in our genetic screen, Rpn11, Doa4 and Ubp14, also display different roles in CytoQC and ERQC (Figure 1, S3 and Amerik et al., 1997; Swaminathan et al., 1999). These differences in DUb functions arise from their diverse subcellular localization and domain organization. Rpn11 is situated at the proteasome but closer to the entry of the catalytic chamber than Ubp6 (Aufderheide et al., 2015; Bashore et al., 2015). Therefore, the *rpn11*^*S119F*^ hypomorphic mutation likely causes a defect in substrate translocation into the proteasome chamber (Verma et al., 2002; Yao and Cohen, 2002), which slows both CytoQC and ERQC and is not rescued by ubiquitin overexpression (Figure S3A and E). Doa4 is physically associated with endosomes by its N-terminal segment (Figure S2 and Amerik et al., 2006) and its deletion results in accumulation of small ubiquitin conjugates (Amerik et al., 2000; Swaminathan et al., 1999). Ubp14 contains zinc finger (ZF) domains which recognize the C-terminus of unanchored polyubiquitin chains to induce an increase in DUb activity (Figure S2 and Amerik et al., 1997; Reyes-Turcu et al., 2006). Because neither *Δdoa4* nor *Δubp14* causes ubiquitin deficiency as severe as *Δubp6*, the accumulation of small ubiquitin conjugates or unanchored polyubiquitin in these strains could be responsible for the decelerated CytoQC (Figure 1A and Amerik et al., 2006; Amerik et al., 1997; Amerik et al., 2000; Swaminathan et al., 1999). However, why they leave ERQC unaffected remains to be explored (Figure 1B).

In conclusion, it is now clear that deletions of individual DUbs do not accelerate QC in *S. cerevisiae*. On the contrary, DUbs such as Ubp6 and Ubp3 promote different QC pathways by distinct mechanisms including ubiquitin recycling and the maintenance of vesicle transport. Further investigation into these diverse mechanisms will aid in our understanding of how CytoQC and ERQC are organized to efficiently clear aberrant proteins.

## Materials and methods

### Euploid yeast strains and culture

Strains used in the gene deletion screen were derived from *S. cerevisiae BY4742* (*s288c his3Δ1 leu2Δ0 lys2Δ0 ura3Δ0*, YSC1049 from Dharmacon). *rpn11*^*S119F*^ and *sec12-4* were derived from *W303* (*leu2-3,112 trp1-1 can1-100 ura3-1 ade2-1 his3-11,15*). Others were derived from *RLY2626* (*s228c ura3 his3 trp1 leu2 LYS2*) (Pavelka et al., 2010). Gene deletion strains were generated by PCR-based gene knock-out (Gietz and Schiestl, 2007; Longtine et al., 1998). Euploid yeast strains were maintained by typical methods (Sherman, 2002), and for experiments, cultured in synthetic media at 30°C to mid-exponential phase (A_600_ ≈ 0.7). A list of *S. cerevisiae* strains used in this study can be found in Table S1.

### Retrieval and generation of DUb mutants

We retrieved the deletion strains of non-essential DUbs, except for Ubp10, from a deletion library sold by Dharmacon. Their identities were re-confirmed by PCR genotyping. *Δubp10* was not provided by the deletion library so we generated this mutant on our own. For Rpn11, which is essential for cell growth, we acquired from a genetic selection for CytoQC components an *rpn11*^*S119F*^ mutant, which is reduced in its Zn^2+^-coordinating ability required for deubiquitination by this metalloprotease (Maytal-Kivity et al., 2002; Tran et al., 2003 and our submitted manuscript).

### Aneuploid yeast strains and culture

*dis XIII* and *dis I, VIII* aneuploid strains are kind gifts from G. Rancati and R. Li (Pavelka et al., 2010). These strains are derivatives of *RLY2626*. Aneuploid strains were always maintained at 25°C. The ploidy of all aneuploid strains was verified by qPCR karyotyping (below). When aneuploid cells were cultured at 30°C for pulse-chase (Figure 3), an aliquot of the same culture was also karyotyped.

### qPCR karyotyping

*S. cerevisiae* cells of the exponential phase were diluted to A_600_ = 0.3. Of the normalized culture, 300 μL was taken and cells were washed with phosphate buffered saline (PBS, pH = 7.5). Then cell walls were digested by 14 mg/mL of zymolyase 20T (US Biologicals Z1000) in 21.5 μL of PBS plus 2.3 mM of DTT. Afterwards, the genomic DNA was released by boiling at 100°C for 5 min. 0.5 μL of the cell lysate was used in qPCR, conducted using reagents and protocol from the QuantiNova SYBR Green PCR Kit (Qiagen 204141). Primers for karyotyping were published previously (Pavelka et al., 2010). The variation of chromosome copy numbers at the population level was kept within ± 0.2 for disomic chromosomes.

### Substrates and plasmids

Substrates of the UPS examined in this study (Figure S1) were hosted on centromeric plasmids. Among them, KWW is HA-tagged at the C-terminus of its KHN domain and Deg1-Ura3 is not tagged. Other proteins are HA-tagged at their C-termini. A list of plasmids used in this study are shown in Table S2. All insertions on plasmids have been validated by sequencing.

### Antibodies

To immunoprecipitate HA-tagged proteins and to detect HA-tagged proteins in immunoblotting, the anti-HA monoclonal mouse antibody HA.11 (BioLegend 901501) was routinely used. Other antibodies, antisera and affinity matrix used in this study are: anti-Ura3 rabbit serum (raised in lab), anti-Ub monoclonal mouse antibody Ubi-1 (invitrogen 13-1600), anti-Pgk1 monoclonal mouse antibody (invitrogen 459250), anti-CPY rabbit serum (kind gift from R. Gilmore), and anti-HA affinity matrix (Roche 11815016001).

### Metabolic ^35^S labelling and pulse-chase

*S. cerevisiae* cells of the mid-exponential phase were concentrated 5 times in fresh media and allowed 30 min to adapt. Cells were then labelled by adding the EXPRE^35^S^35^S Protein Labeling Mix (PerkinElmer EasyTagTM NEG772) at a ratio of 9 μL per mL of the concentrated culture. After 5 or 10 min, pulse-labelling was quenched by adding 12.5 μL of chase media (200 mM methionine, 200 mM cysteine) for each mL of the culture. Then, 1 mL of the concentrated culture was aliquoted at different time-points post-labelling and all cellular activities in the aliquot were immediately killed by adding trichloroacetic acid (TCA) to 10% (v/v). After protein extraction and the immunoprecipitation of substrates (see below), samples were fractionated by SDS-PAGE. Gels were dried and exposed to storage phosphor screens (Kodak SO230, Fuji BAS-IP SR 2025). Finally, the screens were scanned by a Typhoon 9200 Scanner (GE Healthcare) and analyzed in ImageQuant TL.

### Protein (denatured) extraction

Cells killed by 10% TCA were subsequently lysed by bead beating. Then, proteins were precipitated by centrifugation (> 18000 g, 15 min at 4°C) and for each mL of yeast culture in exponential phase, dissolved in 16 - 35 μL of TCA resuspension buffer (3% SDS [w/v], 100 mM Tris pH = 9.0, 3 mM DTT) by boiling and vortexing at 100°C. For ubiquitination assay (below), DTT was omitted from TCA resuspension buffer to extract proteins under non-reducing condition.

### Immunoprecipitation (IP)

50 μL of protein extract (in TCA resuspension buffer) was mixed with 700 μL of IP solution II (20 mM Tris pH = 7.5, 150 mM NaCl, 1% [w/v] Triton X-100, 0.02% [w/v] NaN_3_), 6 μL of 100 mM PMSF, 1 μL of protease inhibitor cocktail (Sigma P8215) and 1 - 5 μL of antibody solution. The mixture was incubated for 1 h under 4°C. After removing insoluble materials by centrifugation (> 18000 g, 20 min at 4°C), the supernatant was mixed with 30 μL of protein A Sepharose (Sigma P3391) and rotated for 2 h at 4°C. Protein A beads were then washed 3 times with IP solution I (IP solution II + 0.2% [w/v] SDS) and 2 times with PBS. Finally, proteins were eluted into ∼ 25 μL of PBS plus 10 μL of 4x Laemmli buffer (125 mM Tris-HCl, pH = 6.8, 4% [w/v] SDS, 50% [v/v] glycerol, 0.2 mg/mL bromophenol blue, 5% [v/v] β-mercaptoethanol) by boiling and vortexing at 100°C.

### Immunoblotting (IB)

Nitrocellulose membranes (BIO-RAD 1620213 or 1704159) were used for the electroblotting of proteasomal substrates. PVDF (BIO-RAD 1704156) was used for the blotting of free Ub and was autoclaved in water after blotting (Swerdlow et al., 1986). After blocking in Odyssey Blocking Buffer (PBS, LI-COR 927), membranes were incubated sequentially with primary and secondary antibodies in Odyssey Blocking Buffer mixed with equal volume of PBS and 0.1% (v/v) of Tween 20. After each incubation, membranes were washed in PBS plus 0.1% (v/v) of Tween 20. Tween 20 was removed by rinsing in PBS before detecting the fluorescence of secondary antibodies using a LI-COR Odyssey Classic Scanner. The fluorescence of protein bands was quantified by Odyssey Application Software while the ubiquitination profiles were quantified in ImageQuant TL by 1D Gel Analysis.

### Ubiquitination assay

Proteins were extracted under non-reducing condition to preserve unconventional ubiquitination on cysteine residues (Baldridge and Rapoport, 2016). Up to 85 μL of the protein extract, normalized to contain equal amounts of un-modified substrates, was mixed with 50 μL of anti-HA affinity matrix, 1200 μL of IP solution II, 1.8 μL of PIC and 10.5 μL of PMSF to immunoprecipitate HA-tagged proteins. Products of IP were fractionated by non-reducing SDS-PAGE and electroblotted (4°C overnight) onto nitrocellulose membranes. The blots were autoclaved to better expose the antigen (Swerdlow et al., 1986) and ubiquitinated species were detected by immunoblotting against ubiquitin (weak fluorescent signal). The non-ubiquitinated species was subsequently detected by blotting against HA (strong fluorescent signal).

## Author contributions

H.W. performed the experiments and analysis. D.T.W.N., H.W., P.T.M. and I.C. conceived the project. H.W., P.T.M. and I.C. wrote the article.

## Acknowledgements

We thank P. Walter and R. Gilmore for gifts of antibodies and G. Rancati and R. Li for sharing aneuploid strains. We are grateful to K. Larrimore for teaching how to handle aneuploids, Y. Liu and N. Bedford for technical support and S.N. Chan, S. Zhang, C. Xu, R. Li, M. Wenk and G. Jedd for constructive discussions. The benchwork of this project was conducted in Temasek Lifesciences Laboratory (TLL) and generously sponsored by TLL core funding. H.W. was a recipient of Graduate Research Scholarship from National University of Singapore and Research Assistantship from P.T.M. lab.

## Abbreviations

*chr*: *chromosome*
CPY: carboxypeptidase Y
CytoQC: cytosolic (protein) quality control
*dis*: *disomic*
DTT: dithiothreitol DUb: deubiquitinase
ER: endoplasmic reticulum
ERQC: ER (protein) quality control
GFP: green fluorescent protein
HA: hemagglutinin tag
IP: immunoprecipitation
PAGE: polyacrylamide gel electrophoresis
PBS: phosphate-buffered saline
PIC: protease inhibitor cocktail
PMSF: phenylmethylsulfonyl fluoride
PrA: proteinase A
*pUB*: *pRS313* centromeric plasmid expressing ubiquitin from *TDH3* promoter
*pUB*^*K63R*^: *pRS424* 2μ plasmid expressing ubiquitin from *PRC1* promoter
QC: (protein) quality control
qPCR: quantitative PCR
SDS: sodium dodecyl sulfate
ss: signal sequence
TCA: trichloroacetic acid
Ub: ubiquitin
UPS: ubiquitin-proteasome system
WT: wild-type

---

**Figure S1.**
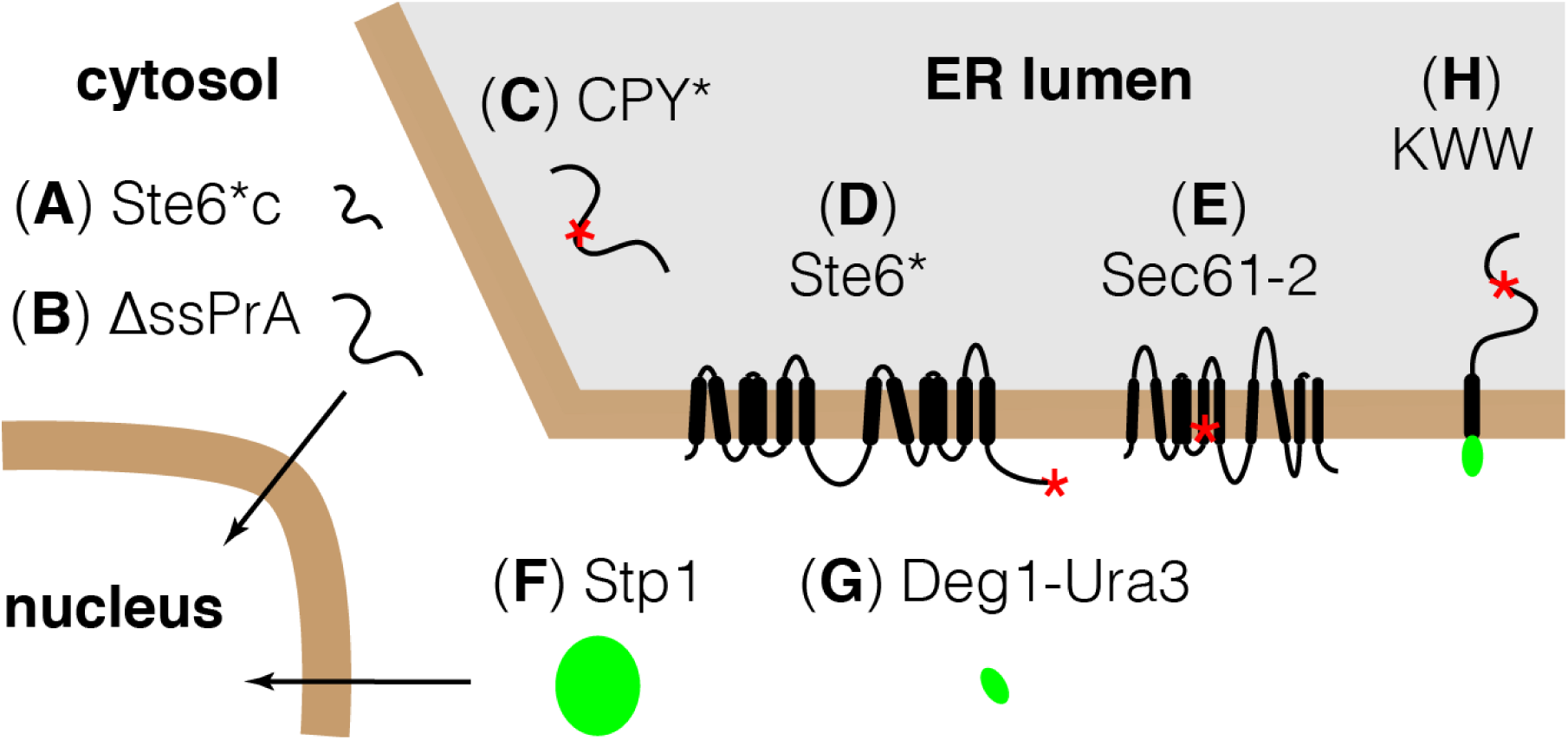
Substrates used in this study. These substrates were selected to cover the canonical pathways of QC and the degradation of folded proteins via the UPS. (**A and B**) CytoQC substrates: Ste6*c and ΔssPrA (Prasad et al., 2010, Prasad et al., 2012). (**C – E and H**) ERQC substrates: (C and H) ERQC-L substrates CPY* and KWW; (D) ERQC-C substrate Ste6*; (E) ERQC-M substrate Sec61-2 (Vashit and Ng, 2004). (**F and G**) Folded UPS substrate Stp1 and Deg1-Ura3 (Gowda et al., 2013; Pfirrmann et al., 2010; Johnson et al., 2010). For each substrate, a schematic diagram is provided. Black vertical cylinder (**|**): transmembrane helix; black curve (⋂): loop region; red asterisk (*): mutation/misfolded site; green oval (●): folded domain; black solid arrow line (→): translocation of substrate. Diagrams are not to scale.

**Figure S2.**
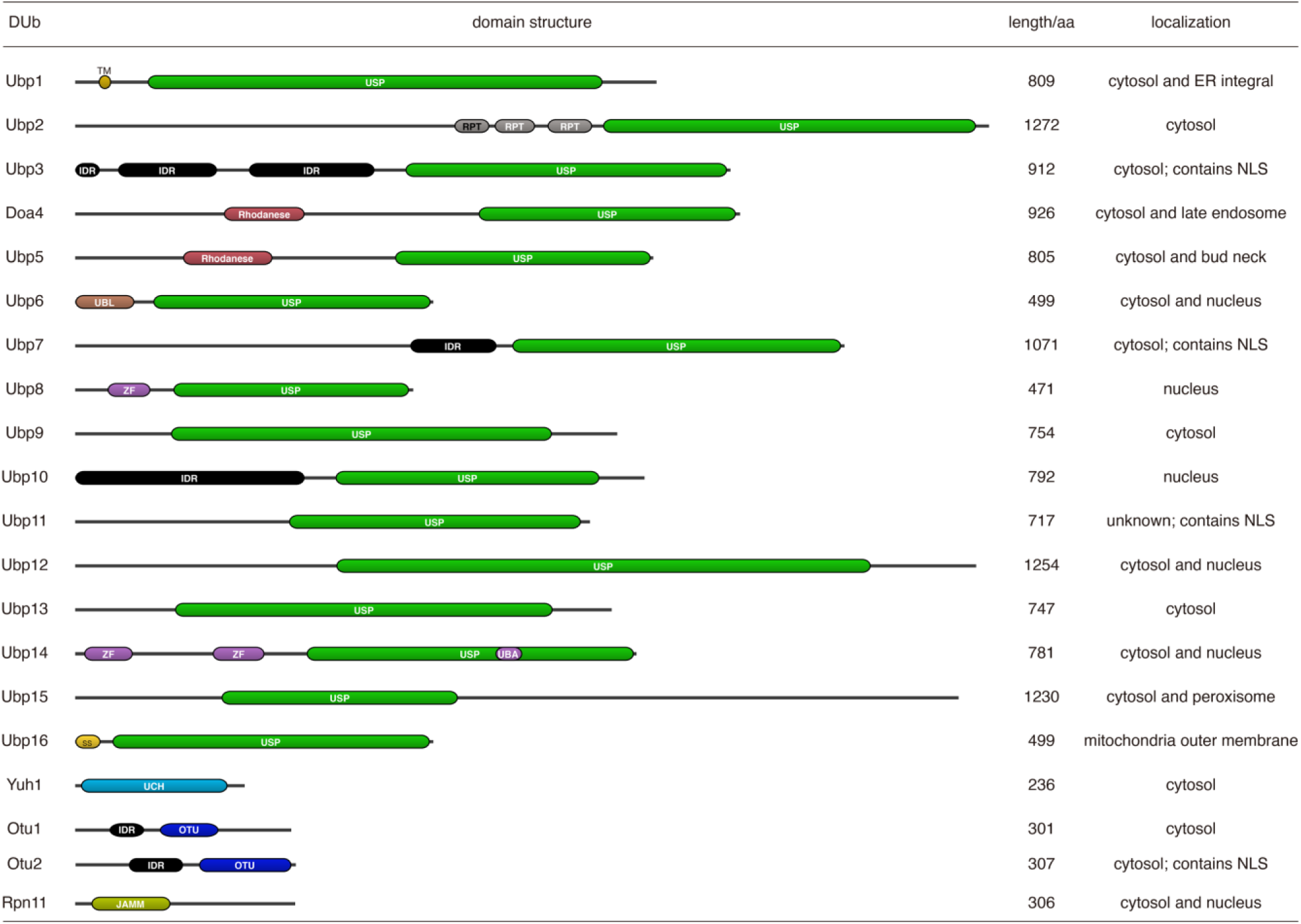
List of yeast DUbs. The catalytic domains of these DUbs fall into 4 families: ubiquitin-specific protease (USP), ubiquitin C-terminal hydrolase (UCH), ovarian tumor (OTU) and JAB1/MPN/Mov34 metalloenzyme (JAMM). Other domains/regions found in DUbs are: transmembrane (TM), repeats (RPT), disordered region (IDR), rhodanese homology domain (rhodanese), ubiquitin-like (UBL) domain, zinc finger (ZF), ubiquitin-associated (UBA) domain and signal sequence (ss). NLS: Nuclear Localization Signal. The domain structures of DUbs were identified in Pfam and rendered in DoMosaics (El-Gebali et al., 2019; Moore et al., 2014). NLS was predicted by NLS Mapper using default settings (Kosugi et al., 2009).

**Figure S3.**
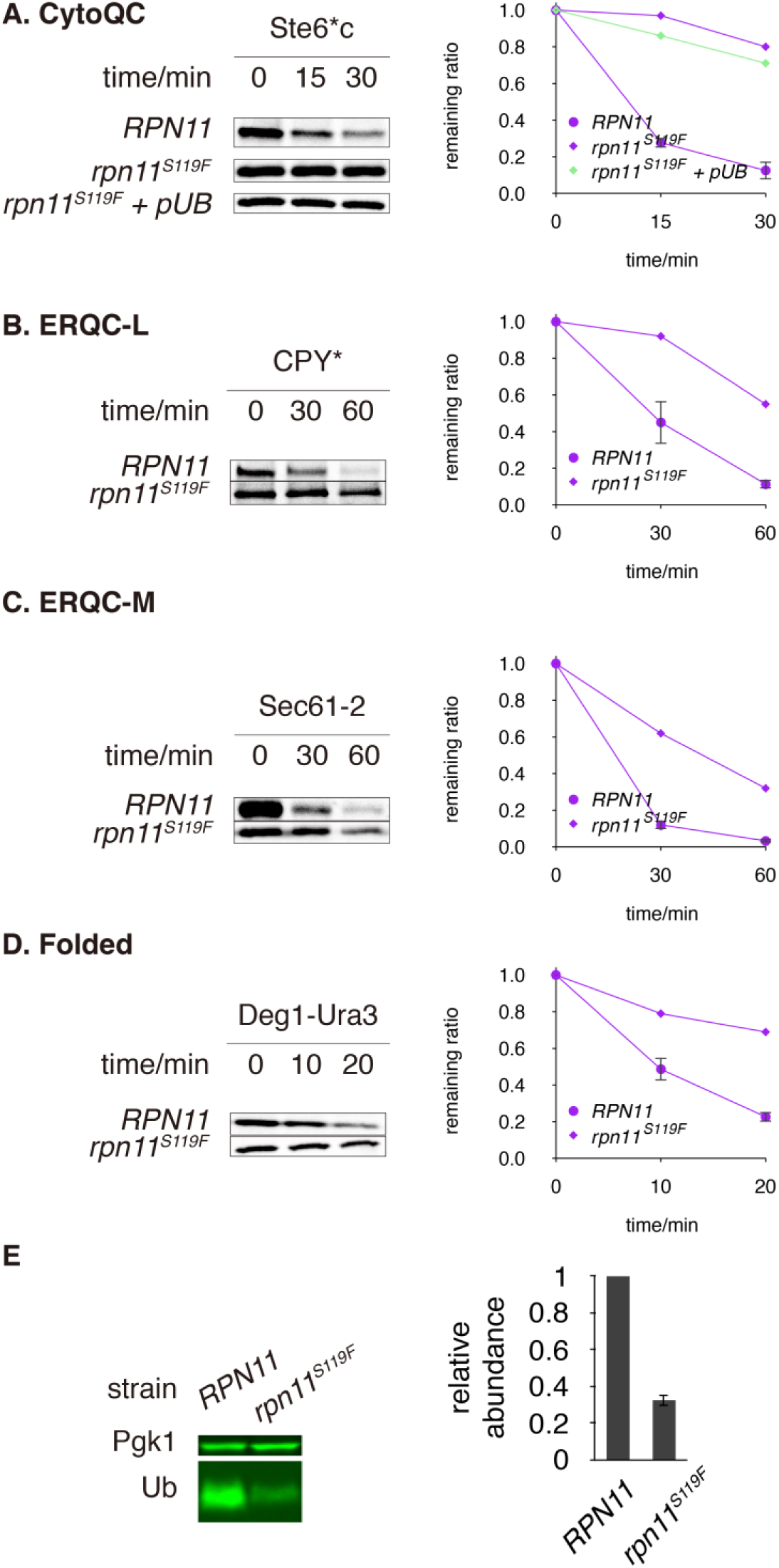
*rpn11* mutation at its active site (*rpn11*^*S119F*^) impairs all degradation pathways via the UPS. (**A**) Degradation of Ste6*c in *RPN11* and *rpn11*^*S119F*^ (*W303* background) in the absence or presence of ubiquitin overexpression. (**B-D**) Degradation of CPY*, Sec61-2 and Deg1-Ura3 in *RPN11* and *rpn11*^*S119F*^. Substrates were pulsed-chased as in Figure 2. Data was processed as described in Figure 2. (**E**) Free ubiquitin abundance in *RPN11* and *rpn11*^*S119F*^, assayed as in Figure 4C.

**Figure S4.**
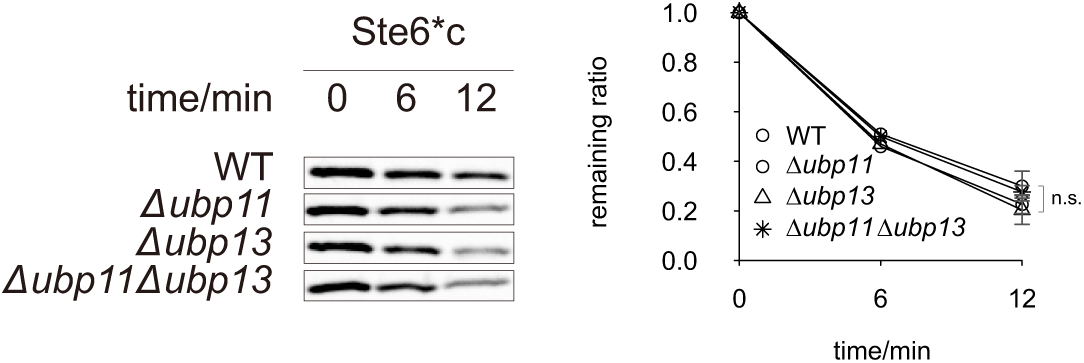
Degradation of Ste6*c in *Δubp11, Δubp13* and *Δubp11Δubp13*, assayed by pulse-chase as in Figure 2.

**Figure S5.**
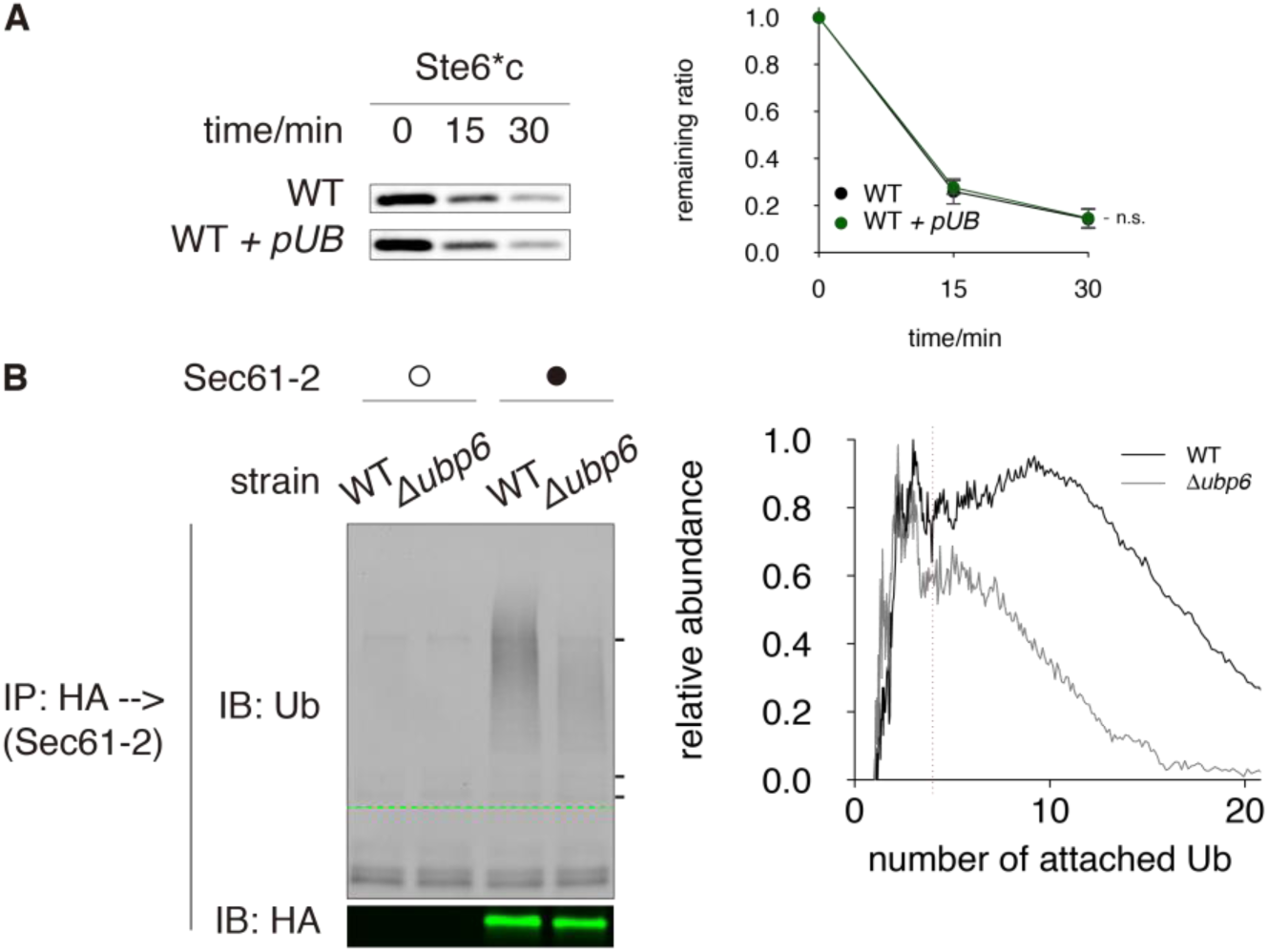
Supplemental to Ubp6 mechanism. (**A**) Degradation of Ste6*c in WT and WT + *pUB*. (**B**) Ubiquitination of Sec61-2, a misfolded ER membrane protein (with lesion in membrane). Products of immunoprecipitation from strains without harboring substrates were run in adjacent lanes as control. Empty circle: vector control; Filled circle: Sec61-2 expressed.

**Figure S6.**
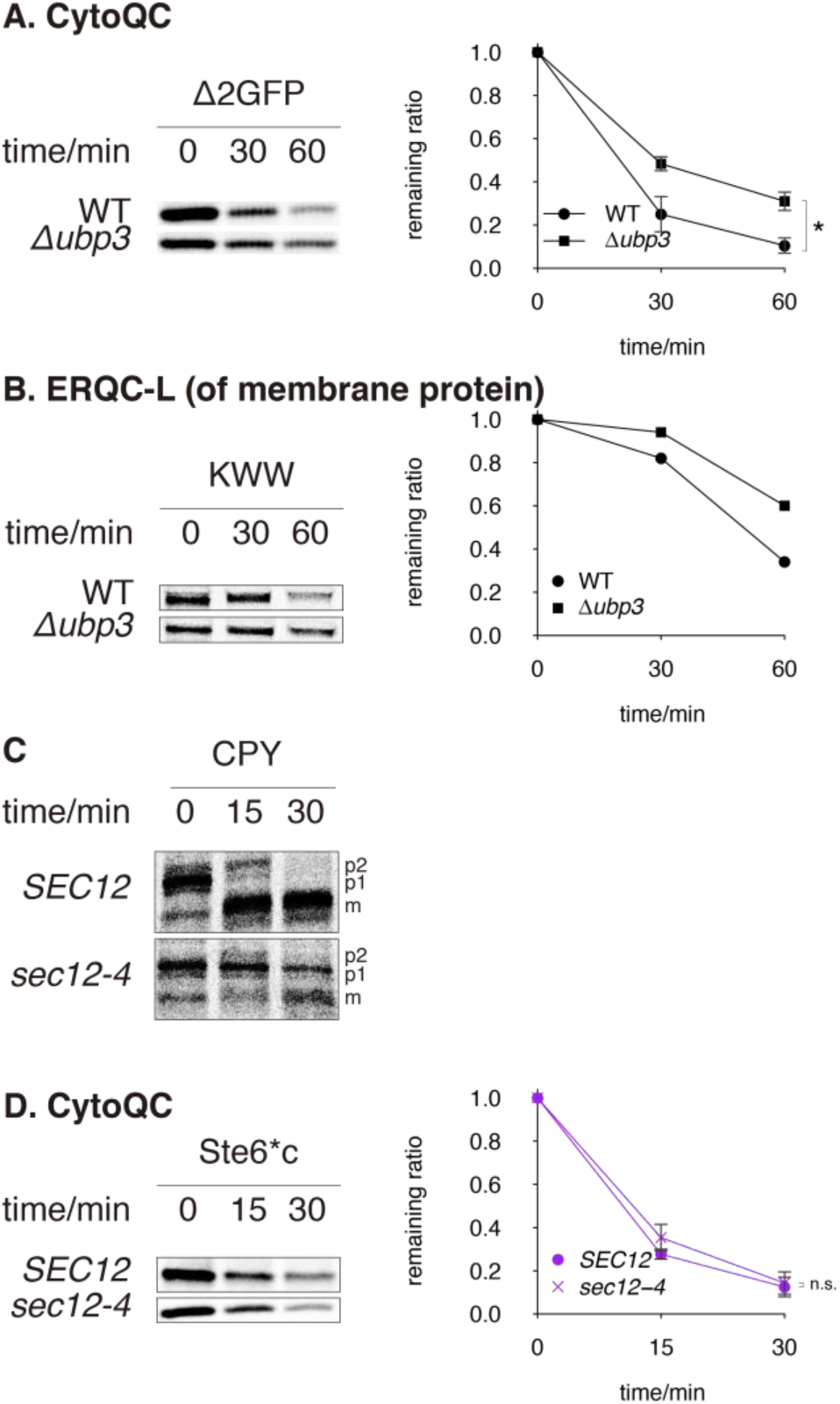
Supplemental to Ubp3 mechanism. (**A and B**) Degradation of Δ2GFP and KWW in *Δubp3* versus WT, assayed by pulse-chase as in Figure 2. (**C**) Maturation of CPY in *sec12-4* (of W303 background) and isogenic WT (*SEC12*). Cells were cultured and pulse-chased at 30°C. (**D**) Degradation of Ste6*c in *sec12-4* (of W303 background) and isogenic WT (*SEC12*). Cells were cultured and pulse-chased at 30°C, the semi-permissive temperature of *sec12-4* (panel C).

**Table S1.**
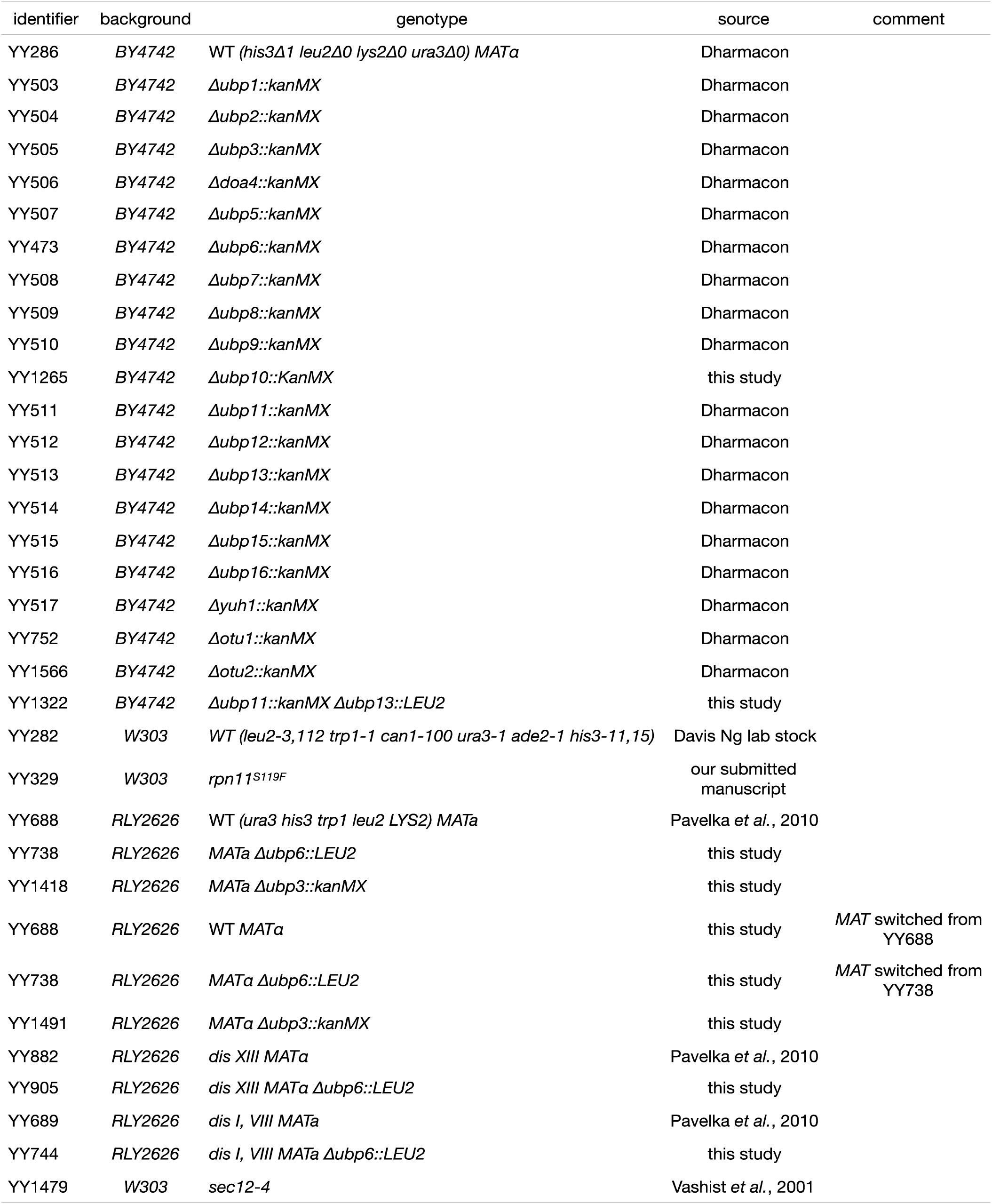
Yeast strains. *kanMX* is a gene cassette that enables yeast to grow with G418 (geneticin).

**Table S2.**
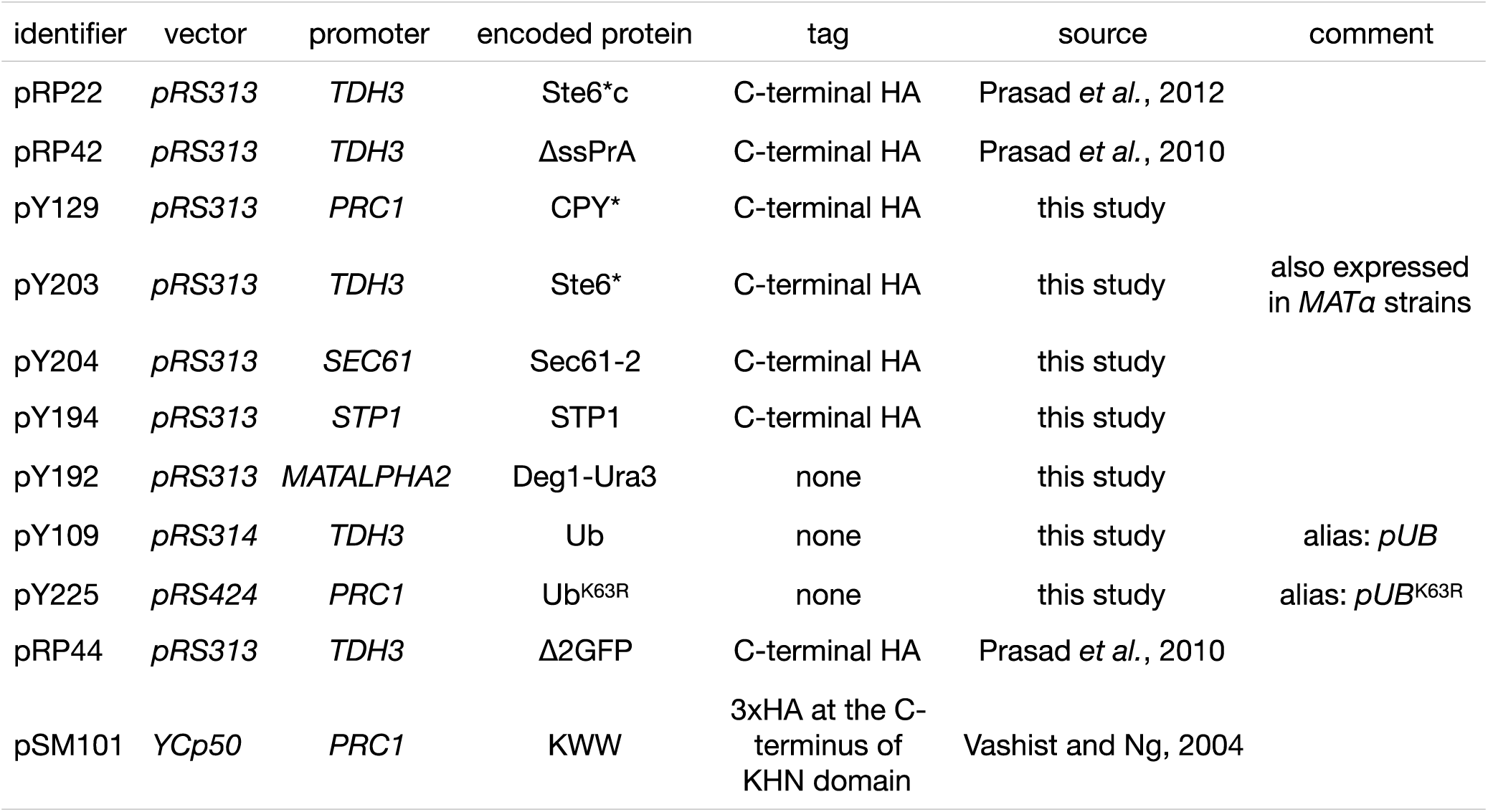
Plasmids. *pRS313, pRS314* and *YCp50* are centromeric vectors while *pRS424* is a 2μ vector (Rose et al., 1987; Sikorski and Hieter, 1989). *TDH3* and *PRC1* promoters are strong and moderate constitutive promoters, respectively. All plasmids harbor *ACT1* terminator downstream the genes expressed. Plasmid maps and sequences are available on request.

## References

Amerik, A., N. Sindhi, and M. Hochstrasser. 2006. A conserved late endosome-targeting signal required for Doa4 deubiquitylating enzyme function. The Journal of cell biology. 175:825–835.

Amerik, A., S. Swaminathan, B.A. Krantz, K.D. Wilkinson, and M. Hochstrasser. 1997. In vivo disassembly of free polyubiquitin chains by yeast Ubp14 modulates rates of protein degradation by the proteasome. The EMBO journal. 16:4826–4838.

Amerik, A.Y., J. Nowak, S. Swaminathan, and M. Hochstrasser. 2000. The Doa4 deubiquitinating enzyme is functionally linked to the vacuolar protein-sorting and endocytic pathways. Molecular biology of the cell. 11:3365–3380.

Ast, T., N. Aviram, S.G. Chuartzman, and M. Schuldiner. 2014. A cytosolic degradation pathway, prERAD, monitors pre-inserted secretory pathway proteins. J Cell Sci. 127:3017–3023.

Aufderheide, A., F. Beck, F. Stengel, M. Hartwig, A. Schweitzer, G. Pfeifer, A.L. Goldberg, E. Sakata, W. Baumeister, and F. Forster. 2015. Structural characterization of the interaction of Ubp6 with the 26S proteasome. Proceedings of the National Academy of Sciences of the United States of America. 112:8626–8631.

Baldridge, R.D., and T.A. Rapoport. 2016. Autoubiquitination of the Hrd1 Ligase Triggers Protein Retrotranslocation in ERAD. Cell. 166:394–407.

Bashore, C., C.M. Dambacher, E.A. Goodall, M.E. Matyskiela, G.C. Lander, and A. Martin. 2015. Ubp6 deubiquitinase controls conformational dynamics and substrate degradation of the 26S proteasome. Nature structural & molecular biology. 22:712–719.

Blount, J.R., A.A. Burr, A. Denuc, G. Marfany, and S.V. Todi. 2012. Ubiquitin-specific protease 25 functions in Endoplasmic Reticulum-associated degradation. PloS one. 7:e36542.

Boselli, M., B.H. Lee, J. Robert, M.A. Prado, S.W. Min, C. Cheng, M.C. Silva, C. Seong, S. Elsasser, K.M. Hatle, T.C. Gahman, S.P. Gygi, S.J. Haggarty, L. Gan, R.W. King, and D. Finley. 2017. An inhibitor of the proteasomal deubiquitinating enzyme USP14 induces tau elimination in cultured neurons. The Journal of biological chemistry. 292:19209–19225.

Chernova, T.A., K.D. Allen, L.M. Wesoloski, J.R. Shanks, Y.O. Chernoff, and K.D. Wilkinson. 2003. Pleiotropic effects of Ubp6 loss on drug sensitivities and yeast prion are due to depletion of the free ubiquitin pool. The Journal of biological chemistry. 278:52102–52115.

Choe, Y.J., S.H. Park, T. Hassemer, R. Korner, L. Vincenz-Donnelly, M. Hayer-Hartl, and F.U. Hartl. 2016. Failure of RQC machinery causes protein aggregation and proteotoxic stress. Nature. 531:191–195.

Cohen, M., F. Stutz, N. Belgareh, R. Haguenauer-Tsapis, and C. Dargemont. 2003a. Ubp3 requires a cofactor, Bre5, to specifically de-ubiquitinate the COPII protein, Sec23. Nature cell biology. 5:661–667.

Cohen, M., F. Stutz, and C. Dargemont. 2003b. Deubiquitination, a new player in Golgi to endoplasmic reticulum retrograde transport. The Journal of biological chemistry. 278:51989–51992.

Dephoure, N., S. Hwang, C. O’Sullivan, S.E. Dodgson, S.P. Gygi, A. Amon, and E.M. Torres. 2014. Quantitative proteomic analysis reveals posttranslational responses to aneuploidy in yeast. eLife. 3:e03023.

El-Gebali, S., J. Mistry, A. Bateman, S.R. Eddy, A. Luciani, S.C. Potter, M. Qureshi, L.J. Richardson, G.A. Salazar, A. Smart, E.L.L. Sonnhammer, L. Hirsh, L. Paladin, D. Piovesan, S.C.E. Tosatto, and R.D. Finn. 2019. The Pfam protein families database in 2019. Nucleic acids research. 47:D427–d432.

Fang, N.N., G.T. Chan, M. Zhu, S.A. Comyn, A. Persaud, R.J. Deshaies, D. Rotin, J. Gsponer, and T. Mayor. 2014. Rsp5/Nedd4 is the main ubiquitin ligase that targets cytosolic misfolded proteins following heat stress. Nature cell biology. 16:1227–1237.

Fang, N.N., M. Zhu, A. Rose, K.P. Wu, and T. Mayor. 2016. Deubiquitinase activity is required for the proteasomal degradation of misfolded cytosolic proteins upon heat-stress. Nature communications. 7:12907.

Finley, D., H.D. Ulrich, T. Sommer, and P. Kaiser. 2012. The ubiquitin-proteasome system of Saccharomyces cerevisiae. Genetics. 192:319–360.

Gietz, R.D., and R.H. Schiestl. 2007. High-efficiency yeast transformation using the LiAc/SS carrier DNA/PEG method. Nature protocols. 2:31–34.

Gowda, N.K., G. Kandasamy, M.S. Froehlich, R.J. Dohmen, and C. Andreasson. 2013. Hsp70 nucleotide exchange factor Fes1 is essential for ubiquitin-dependent degradation of misfolded cytosolic proteins. Proceedings of the National Academy of Sciences of the United States of America. 110:5975–5980.

Guerriero, C.J., K.R. Reutter, A. Augustine, and J.L. Brodsky. 2016. The degradation requirements for topologically distinct quality control substrates in the yeast endoplasmic reticulum. The FASEB Journal. 30:1063.1062-1063.1062.

Guerriero, C.J., K.F. Weiberth, and J.L. Brodsky. 2013. Hsp70 targets a cytoplasmic quality control substrate to the San1p ubiquitin ligase. The Journal of biological chemistry. 288:18506–18520.

Guterman, A., and M.H. Glickman. 2004. Complementary roles for Rpn11 and Ubp6 in deubiquitination and proteolysis by the proteasome. The Journal of biological chemistry. 279:1729–1738.

Hampton, R.Y., and T. Sommer. 2012. Finding the will and the way of ERAD substrate retrotranslocation. Current opinion in cell biology. 24:460–466.

Hanna, J., N.A. Hathaway, Y. Tone, B. Crosas, S. Elsasser, D.S. Kirkpatrick, D.S. Leggett, S.P. Gygi, R.W. King, and D. Finley. 2006. Deubiquitinating enzyme Ubp6 functions noncatalytically to delay proteasomal degradation. Cell. 127:99–111.

Hanna, J., D.S. Leggett, and D. Finley. 2003. Ubiquitin depletion as a key mediator of toxicity by translational inhibitors. Molecular and cellular biology. 23:9251–9261.

Heck, J.W., S.K. Cheung, and R.Y. Hampton. 2010. Cytoplasmic protein quality control degradation mediated by parallel actions of the E3 ubiquitin ligases Ubr1 and San1. Proceedings of the National Academy of Sciences of the United States of America. 107:1106–1111.

Hickey, C.M. 2016. Degradation elements coincide with cofactor binding sites in a short-lived transcription factor. Cellular logistics. 6:e1157664.

Kaushik, S., and A.M. Cuervo. 2015. Proteostasis and aging. Nature medicine. 21:1406–1415.

Kawaguchi, S., C.L. Hsu, and D.T. Ng. 2010. Interplay of substrate retention and export signals in endoplasmic reticulum quality control. PloS one. 5:e15532.

Kim, H.T., K.P. Kim, F. Lledias, A.F. Kisselev, K.M. Scaglione, D. Skowyra, S.P. Gygi, and A.L. Goldberg. 2007. Certain pairs of ubiquitin-conjugating enzymes (E2s) and ubiquitin-protein ligases (E3s) synthesize nondegradable forked ubiquitin chains containing all possible isopeptide linkages. The Journal of biological chemistry. 282:17375–17386.

Klabonski, L., J. Zha, L. Senthilkumar, and T. Gidalevitz. 2016. A Bystander Mechanism Explains the Specific Phenotype of a Broadly Expressed Misfolded Protein. PLoS Genet. 12:e1006450.

Kosugi, S., M. Hasebe, M. Tomita, and H. Yanagawa. 2009. Systematic identification of cell cycle-dependent yeast nucleocytoplasmic shuttling proteins by prediction of composite motifs. Proceedings of the National Academy of Sciences of the United States of America. 106:10171–10176.

Lee, D.H., M.Y. Sherman, and A.L. Goldberg. 1996. Involvement of the molecular chaperone Ydj1 in the ubiquitin-dependent degradation of short-lived and abnormal proteins in Saccharomyces cerevisiae. Molecular and cellular biology. 16:4773–4781.

Leggett, D.S., J. Hanna, A. Borodovsky, B. Crosas, M. Schmidt, R.T. Baker, T. Walz, H. Ploegh, and D. Finley. 2002. Multiple associated proteins regulate proteasome structure and function. Molecular cell. 10:495–507.

Liu, Y., N. Soetandyo, J.G. Lee, L. Liu, Y. Xu, W.M. Clemons, Jr., and Y. Ye. 2014. USP13 antagonizes gp78 to maintain functionality of a chaperone in ER-associated degradation. eLife. 3:e01369.

Longtine, M.S., A. McKenzie, 3rd, D.J. Demarini, N.G. Shah, A. Wach, A. Brachat, P. Philippsen, and J.R. Pringle. 1998. Additional modules for versatile and economical PCR-based gene deletion and modification in Saccharomyces cerevisiae. Yeast (Chichester, England). 14:953–961.

Maytal-Kivity, V., N. Reis, K. Hofmann, and M.H. Glickman. 2002. MPN+, a putative catalytic motif found in a subset of MPN domain proteins from eukaryotes and prokaryotes, is critical for Rpn11 function. BMC biochemistry. 3:28.

Moore, A.D., A. Held, N. Terrapon, J. Weiner, 3rd, and E. Bornberg-Bauer. 2014. DoMosaics: software for domain arrangement visualization and domain-centric analysis of proteins. Bioinformatics (Oxford, England). 30:282–283.

Nakatsukasa, K., and T. Kamura. 2016. Subcellular Fractionation Analysis of the Extraction of Ubiquitinated Polytopic Membrane Substrate during ER-Associated Degradation. PloS one. 11:e0148327.

Nielsen, S.V., E.G. Poulsen, C.A. Rebula, and R. Hartmann-Petersen. 2014. Protein quality control in the nucleus. Biomolecules. 4:646–661.

Oling, D., F. Eisele, K. Kvint, and T. Nystrom. 2014. Opposing roles of Ubp3-dependent deubiquitination regulate replicative life span and heat resistance. The EMBO journal. 33:747–761.

Ozkaynak, E., D. Finley, M.J. Solomon, and A. Varshavsky. 1987. The yeast ubiquitin genes: a family of natural gene fusions. The EMBO journal. 6:1429–1439.

Park, S.H., N. Bolender, F. Eisele, Z. Kostova, J. Takeuchi, P. Coffino, and D.H. Wolf. 2007. The cytoplasmic Hsp70 chaperone machinery subjects misfolded and endoplasmic reticulum import-incompetent proteins to degradation via the ubiquitin-proteasome system. Molecular biology of the cell. 18:153–165.

Park, S.H., Y. Kukushkin, R. Gupta, T. Chen, A. Konagai, M.S. Hipp, M. Hayer-Hartl, and F.U. Hartl. 2013. PolyQ proteins interfere with nuclear degradation of cytosolic proteins by sequestering the Sis1p chaperone. Cell. 154:134–145.

Pavelka, N., G. Rancati, J. Zhu, W.D. Bradford, A. Saraf, L. Florens, B.W. Sanderson, G.L. Hattem, and R. Li. 2010. Aneuploidy confers quantitative proteome changes and phenotypic variation in budding yeast. Nature. 468:321–325.

Prasad, R., S. Kawaguchi, and D.T. Ng. 2010. A nucleus-based quality control mechanism for cytosolic proteins. Molecular biology of the cell. 21:2117–2127.

Prasad, R., S. Kawaguchi, and D.T. Ng. 2012. Biosynthetic mode can determine the mechanism of protein quality control. Biochemical and biophysical research communications. 425:689–695.

Prasad, R., C. Xu, and D.T.W. Ng. 2018. Hsp40/70/110 chaperones adapt nuclear protein quality control to serve cytosolic clients. The Journal of cell biology. 217:2019–2032.

Reyes-Turcu, F.E., J.R. Horton, J.E. Mullally, A. Heroux, X. Cheng, and K.D. Wilkinson. 2006. The ubiquitin binding domain ZnF UBP recognizes the C-terminal diglycine motif of unanchored ubiquitin. Cell. 124:1197–1208.

Reyes-Turcu, F.E., K.H. Ventii, and K.D. Wilkinson. 2009. Regulation and cellular roles of ubiquitin-specific deubiquitinating enzymes. Annual review of biochemistry. 78:363–397.

Rose, M.D., P. Novick, J.H. Thomas, D. Botstein, and G.R. Fink. 1987. A Saccharomyces cerevisiae genomic plasmid bank based on a centromere-containing shuttle vector. Gene. 60:237–243.

Rumpf, S., and S. Jentsch. 2006. Functional division of substrate processing cofactors of the ubiquitin-selective Cdc48 chaperone. Molecular cell. 21:261–269.

Sherman, F. 2002. Getting started with yeast. In Methods in Enzymology. Vol. 350. 3–41.

Sikorski, R.S., and P. Hieter. 1989. A system of shuttle vectors and yeast host strains designed for efficient manipulation of DNA in Saccharomyces cerevisiae. Genetics. 122:19–27.

Silva, G.M., D. Finley, and C. Vogel. 2015. K63 polyubiquitination is a new modulator of the oxidative stress response. Nature structural & molecular biology. 22:116–123.

Swaminathan, S., A.Y. Amerik, and M. Hochstrasser. 1999. The Doa4 deubiquitinating enzyme is required for ubiquitin homeostasis in yeast. Molecular biology of the cell. 10:2583–2594.

Swerdlow, P.S., D. Finley, and A. Varshavsky. 1986. Enhancement of immunoblot sensitivity by heating of hydrated filters. Analytical biochemistry. 156:147–153.

Taxis, C., F. Vogel, and D.H. Wolf. 2002. ER-golgi traffic is a prerequisite for efficient ER degradation. Molecular biology of the cell. 13:1806–1818.

Tobias, J.W., and A. Varshavsky. 1991. Cloning and functional analysis of the ubiquitin-specific protease gene UBP1 of Saccharomyces cerevisiae. The Journal of biological chemistry. 266:12021–12028.

Torres, E.M., N. Dephoure, A. Panneerselvam, C.M. Tucker, C.A. Whittaker, S.P. Gygi, M.J. Dunham, and A. Amon. 2010. Identification of aneuploidy-tolerating mutations. Cell. 143:71–83.

Tran, A. 2019. The N-end rule pathway and Ubr1 enforce protein compartmentalization via P2-encoded cellular location signals. J Cell Sci. 132.

Tran, H.J., M.D. Allen, J. Lowe, and M. Bycroft. 2003. Structure of the Jab1/MPN domain and its implications for proteasome function. Biochemistry. 42:11460–11465.

Tumusiime, S., C. Zhang, M.S. Overstreet, and Z. Liu. 2011. Differential regulation of transcription factors Stp1 and Stp2 in the Ssy1-Ptr3-Ssy5 amino acid sensing pathway. The Journal of biological chemistry. 286:4620–4631.

Vashist, S., W. Kim, W.J. Belden, E.D. Spear, C. Barlowe, and D.T. Ng. 2001. Distinct retrieval and retention mechanisms are required for the quality control of endoplasmic reticulum protein folding. The Journal of cell biology. 155:355–368.

Vashist, S., and D.T. Ng. 2004. Misfolded proteins are sorted by a sequential checkpoint mechanism of ER quality control. The Journal of cell biology. 165:41–52.

Verma, R., L. Aravind, R. Oania, W.H. McDonald, J.R. Yates, 3rd, E.V. Koonin, and R.J. Deshaies. 2002. Role of Rpn11 metalloprotease in deubiquitination and degradation by the 26S proteasome. Science (New York, N.Y.). 298:611–615.

Wang, X., Y.Y. Yu, N. Myers, and T.H. Hansen. 2013. Decoupling the role of ubiquitination for the dislocation versus degradation of major histocompatibility complex (MHC) class I proteins during endoplasmic reticulum-associated degradation (ERAD). The Journal of biological chemistry. 288:23295–23306.

Yao, T., and R.E. Cohen. 2002. A cryptic protease couples deubiquitination and degradation by the proteasome. Nature. 419:403–407.

Yu, L., Y. Chen, and S.A. Tooze. 2018. Autophagy pathway: Cellular and molecular mechanisms. Autophagy. 14:207–215.

Zattas, D., J.M. Berk, S.G. Kreft, and M. Hochstrasser. 2016. A Conserved C-terminal Element in the Yeast Doa10 and Human MARCH6 Ubiquitin Ligases Required for Selective Substrate Degradation. The Journal of biological chemistry. 291:12105–12118.

Zhang, Z.R., J.S. Bonifacino, and R.S. Hegde. 2013. Deubiquitinases sharpen substrate discrimination during membrane protein degradation from the ER. Cell. 154:609–622.

